# Calponin 2 harnesses metabolic reprogramming to determine kidney fibrosis

**DOI:** 10.1101/2023.01.03.522608

**Authors:** Yuan Gui, Jianling Tao, Yuanyuan Wang, Zachary Palanza, Yi Qiao, Geneva Hargis, Donald L. Kreutzer, Silvia Liu, Sheldon I. Bastacky, Yanlin Wang, Yanbao Yu, Haiyan Fu, Dong Zhou

## Abstract

In the fibrotic kidneys, the extent of a formed deleterious microenvironment is determined by cellular mechanical forces. This process requires metabolism for energy; however, how cellular mechanics and metabolism are connected remains unclear. Our proteomics revealed that actin filament binding and cell metabolism are the two most dysregulated events in the fibrotic kidneys. As a prominent actin stabilizer, Calponin 2 (CNN2) is predominantly expressed in fibroblasts and pericytes. CNN2 knockdown preserves kidney function and alleviates fibrosis. Global proteomics profiled that CNN2 knockdown enhanced the activities of the key rate-limiting enzymes and regulators of fatty acid oxidation (FAO) in diseased kidneys. Inhibiting carnitine palmitoyltransferase 1α in the FAO pathway results in lipid accumulation and extracellular matrix deposition in the fibrotic kidneys, which were restored after CNN2 knockdown. In patients, increased serum CNN2 levels are correlated with lipid content. Bioinformatics and chromatin immunoprecipitation showed that CNN2 interactor, estrogen receptor 2 (ESR2) binds peroxisome proliferator-activated receptor-α (PPARα) to transcriptionally regulate FAO downstream target genes expression amid kidney fibrosis. *In vitro*, ESR2 knockdown repressed the mRNA levels of PPARα and the key genes in the FAO pathway. Conversely, activation of PPARα reduced CNN2-induced matrix inductions. Our results suggest that balancing cell mechanics and metabolism is crucial to develop therapeutic strategies to halt kidney fibrosis.

## Introduction

Chronic kidney disease (CKD) is estimated to become the 5^th^ leading cause of death worldwide by 2040.^1,2^ As the common consequence of CKD, kidney fibrosis refers to a heterogeneous group of disorders that scar kidneys, most often inevitably and irreversibly.^3^ In fibrosis, cell mechanical forces are essential in controlling fibroblast activation and extracellular matrix (ECM) stiffness, and regulating tubular cell metabolic reprogramming for energy.^4,5^ Nevertheless, mechanotransduction is often ignored in studying CKD.

Mechanical signals are equally important to chemical signals in exerting biological effects. Our proteomics data revealed that actin filament binding is a key event in the fibrotic kidneys. Serving as an actin stabilizer, Calponin (CNN) plays a central role in numerous fundamental biological processes, including cell proliferation, motility, and adhesion.^6,7^ CNN mostly consists of α-helices with hydrogen bond turns. As a binding protein, CNN structure consists of three domains. These domains appear in order of CNN homology, regulatory domain, and the domain containing the CNN repeats. In the human genome, CNN has three isoforms: CNN1, CNN2, and CNN3.^6^ In comparison to CNN1 and CNN3, CNN2 is expressed in more tissues and cell types.^6,8^ Under pathophysiological conditions, CNN2 deletion attenuates inflammatory arthritis,^9^ calcific aortic valve disease,^10^ postoperative peritoneal adhesions,^11^ and tumor metastasis.^12^ However, little is known about how CNN2 influences kidney fibrosis.

Cell mechanics and metabolism are intrinsically intertwined.^13^ After CKD, tubular epithelial cells (TECs), the largest resident cell population in the kidney, can adapt to the fibrotic microenvironment by reprogramming their metabolism to meet the energy consumption requirements and improve CKD.^14,15^ Importantly, this metabolic reprogramming provides energy for actin cytoskeletal rearrangement in response to external forces.^16^ TECs demand high energy which primarily relies on fatty acid oxidation (FAO) and mitochondrial oxidative phosphorylation to generate adenosine triphosphate (ATP) as their energy source.^17^ Under CKD, defective FAO induces intracellular lipid accumulation and ATP depletion. Consequently, cellular lipotoxicity and energy deprivation contribute to TEC dedifferentiation, leading to irreversible kidney fibrosis.^18-20^ Fatty acid (FA) is transported to the kidney mainly by the cluster of differentiation 36 (CD36) or FA transporter 2.^21,22^ It is then catalyzed to fatty acyl-CoA by acyl-CoA synthetase (Acsm) in cells.^23^ Activated FA enters the mitochondrial matrix for β-oxidation mediated by carnitine palmitoyltransferase 1 (CPT1) and 2 (CPT2).^24^ In mitochondria, key rate-limiting enzymes such as CPT1, peroxisomal acyl-CoA oxidase (ACOX), acyl-CoA dehydrogenase medium chain (Acadm) or long chain (Acadl) are required to degrade fatty acyl-CoA into the end-product, acetyl-CoA, by FA β-oxidation.^25^ Lastly, acetyl-CoA enters the tricarboxylic acid (TCA) cycle to generate ATP.^26^ Peroxisome proliferator-activated receptor α (PPARα) transcriptionally controls expression of FAO genes to balance FA uptake and oxidation and determine CKD outcomes.^15,20,27^ Because epithelial-mesenchymal communication is key for controlling CKD progression,^28^ we hypothesized that CNN2 affects the FAO pathway and causes kidney fibrosis. Understanding cellular mechano-metabolic changes will create therapeutic opportunities to treat CKD.

## Methods

Detailed methods are in the Supplementary Methods. We used ARRIVE reporting guidelines for this report.^29^

### Mouse models of CKD

Male Balb/c mice were obtained from the Jackson Laboratory. Mouse CKD models were induced by ischemia-reperfusion injury (IRI), unilateral ureteral obstruction (UUO), and adriamycin (ADR), as described previously.^30,31^ Customized short hairpin RNA (ShRNA) specific for the mouse CNN2 gene was ordered from Qianlong Biotech. In a separate experiment, etomoxir was administrated in mice 2 days before IRI and nephrectomy, respectively. All animal experiments were approved by the Institutional Animal Care and Use Committee at the University of Connecticut, School of Medicine.

### Human Kidney Biopsy Specimens

Human kidney biopsy specimens and non-tumor kidney tissue were obtained from the pathology archive at the University of Pittsburgh Medical Center. All procedures followed ethical standards and were approved by the Institutional Review Board at the University of Pittsburgh, School of Medicine. Demographic data are in Supplementary Table S1.

### Proteomic analysis

Kidney tissues were processed following a shotgun proteomics workflow published previously.^32^ The LC-MS/MS analysis was performed using an Ultimate 3000 nanoLC and Q Exactive mass spectrometer system.

### Human serum metabolomics

The serum samples used in this study came from the investigator-initiated clinical trial (ChiCTR2000028949). All participants signed consent forms so extra samples could be used for academic purposes. Serum metabolomics was performed as we previously reported.^33^ Participant demographic and clinical characteristics are in Supplementary Table S2.

### Determination of blood urea nitrogen and serum creatinine

Blood urea nitrogen (BUN) and serum creatinine (Scr) levels were determined using the QuantiChrom™ Urea and Creatinine assay kits, according to the protocols specified by the manufacturer.

### Quantitative Real-Time Reverse Transcription PCR (qRT-PCR)

Total RNA isolation and qRT-PCR were conducted by procedures described previously.^30^ The mRNA levels of various genes were calculated after normalizing with β-actin or GAPDH. The primer sequences used are in Supplementary Table S3.

### Western Blot Analysis

Protein expression was analyzed by western blot as described previously.^30^ Antibodies used are in Supplementary Table S4.

### Triglyceride and ATP measurement

The levels of serum and kidney triglyceride or kidney ATP content were measured by using the Triglyceride Quantification Kit or the ATP Colorimetric/Fluorometric Assay Kit, according to manufacturers’ instructions.

### Enzyme-linked immunosorbent assay (ELISA)

The human and rat calponin 2 ELISA kits were purchased from MyBioSource, Inc. This assay employs the quantitative sandwich enzyme immunoassay technique.

### Histology, and Oil Red O, immunohistochemical, and immunofluorescence staining

Paraffin-embedded kidney sections and cryosections were prepared by a routine procedure. The sections were stained with Masson Trichrome Staining reagents or Oil Red O staining reagents. Immunostaining was performed according to the established protocol as described previously.^30^ Antibodies used are in Supplementary Table S4.

### Cell culture, small interfering RNA (siRNA) inhibition, and treatment

Normal rat kidney fibroblasts (NRK-49F) and human proximal tubular cells (HK-2) were obtained from the American Type Culture Collection. For conditioned media (CM) collection, NRK-49F were transfected with siCNN2 for 24 hours and then cultured with serum-free media for 24 hours. Serum-starved HK-2 cells were then transfected with ESR2-Dicer siRNA or treated with CM, CNN2 recombinant protein, etomoxir, or fenofibrate.

### Chromatin immunoprecipitation (ChIP) Assay

To analyze interactions between ESR2 and the binding sites in the PPARα gene promoter, a ChIP assay was performed. This assay was conducted according to the manufacturer-specified protocols. Primer sequences used for the ChIP assay are in Supplemental Table S3.

### Statistics

All data were expressed as mean ± SEM if not specified otherwise in the legends. Statistical analysis of the data was performed using GraphPad Prism 9. Results are presented in dot plots. *P* < 0.05 was considered statistically significant.

## Results

### Global proteome and metabolome characterization after CKD

At the genome-wide transcript level, metabolism is a key pathway in metabolic disease-associated CKD patients.^19^ To investigate whether metabolism is also involved in non-diabetic CKD, we used quantitative proteomics to profile the kidney proteome landscape of controls and IRI-induced CKD mice. The principal component analysis (PCA) of our proteomic data classified control and IRI mice according to their genotypes (Supplementary Figure S1a). In total, we identified 6,146 proteins and 1,340 of these had significantly different expression (Permutation FDR 0.05) between control and ischemic kidneys (Figure 1a). Compared to controls, 1,127 and 213 proteins were up- and down-regulated in the ischemic kidneys. Correlations between biological replicates within the same group and the distribution of protein intensity are presented (Supplementary Figure S1b-S1c). Kyoto Encyclopedia of Genes and Genomes (KEGG) enrichment analysis highlighted that metabolic pathways are the most significantly disturbed events (Figure 1b), consistent with transcript-level results.^19^ To further evaluate how metabolic changes influence CKD, we compared sera from healthy adults (HA, n=441) and non-diabetic CKD patients (n=338). Metabolomics were performed on sera as previously described.^33^ We identified 97 differentially expressed serum metabolites between HA and CKD patients (Figure 1c; Supplementary Table S5; Supplementary Figure S1d). Partial least squares (PLS) regression indicates that metabolite distribution patterns are different between HA and CKD patients (Figure 1d). We used these differentially expressed metabolites to predict CKD by employing five machine learning algorithms: linear discriminant analysis, support vector machine, random forest, logistic regression, and PLS. The receiver operating characteristic curves revealed that these metabolites could accurately predict HA *versus* CKD and balance the sensitivity and specificity rates (Youden index) (Figure 1e). Notably, about 47.4% of the metabolites in circulation were associated with glycolipid metabolisms such as glucose-1-phosphate and glycerol-3-galactoside (Figure 1f; Supplementary Figure S1e).

**Figure 1.**
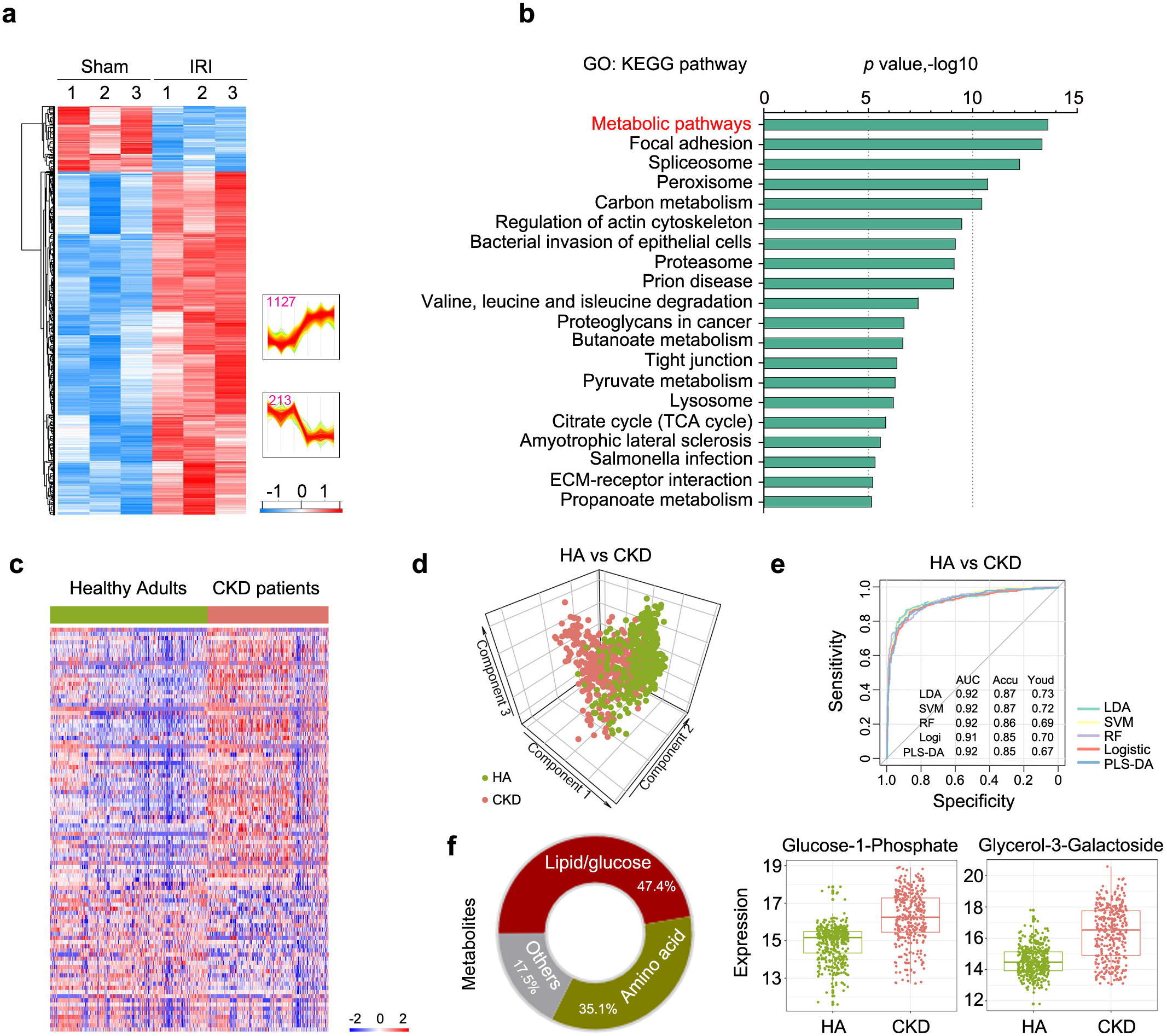
The landscape of proteomes and metabolomes in CKD. (**a**) Hierarchical clustering of the intensity (plotted as z-score) of the proteins identified in the control and ischemic kidneys by quantitative proteomic analysis. Two clusters of proteins with different patterns of abundance profiles were highlighted in the heatmap. (**b**) KEGG pathway enrichment analyses revealed the top 20 activated pathways in the ischemic kidneys after CKD. (**c**) Heatmap of the serum metabolomics data for the comparison between healthy adults and non-diabetic CKD patients. (**d**) Partial least squares discriminant analysis (PLS-DA) based on the differentially expressed metabolites. (**e**) Receiver operating characteristic (ROC) curves for pairwise prediction by five different machine learning methods. Green line for linear discriminant analysis (LDA), yellow line for support vector machine (SVM), purple line for random forest (RF), red line for logistic regression (Logi), and blue line for PLS-DA. AUC: the area under the curve; Accu: accuracy; Youd: Youden index. (**f**) Pie chart showed the ratio of different types of serum metabolites. The dot plots indicated the top two serum metabolites in CKD patients compared to healthy adults.

### Calponin 2 is a key actin regulatory protein that facilitates kidney fibrosis

Besides the disturbance of metabolic pathways, Gene Ontology (GO) analysis of molecular function revealed that actin filament binding plays a key role in facilitating kidney fibrosis (Figure 2a). Because cell mechanics and metabolism are reciprocally regulated,^13^ we screened the significant actin filament proteins in our proteomics database. To identify the most impacted cell mechanics-related proteins for investigation, we excluded unknown proteins and those that have been well studied (Figure 2b). Among the remaining proteins, we identified CNN2 as a protein of interest (Figure 2c). To enhance the reproducibility of our proteomic analyses, we constructed three well-characterized CKD models induced by IRI, UUO, and ADR, respectively. CNN2 mRNA was significantly increased in all three models’ fibrotic kidneys compared to controls (Figure 2d). Western blots demonstrated CNN2 protein was upregulated in fibrotic kidneys (Figure 2e; Supplementary Figure S2, a-c). A separate single nucleus RNA sequencing revealed that kidney fibroblasts and pericytes express CNN2 after IRI (Figure 2f).^34^ Consistently, immunohistochemical staining confirmed that CNN2 was predominantly localized in the fibrotic kidney interstitium (Figure 2g).

**Figure 2:**
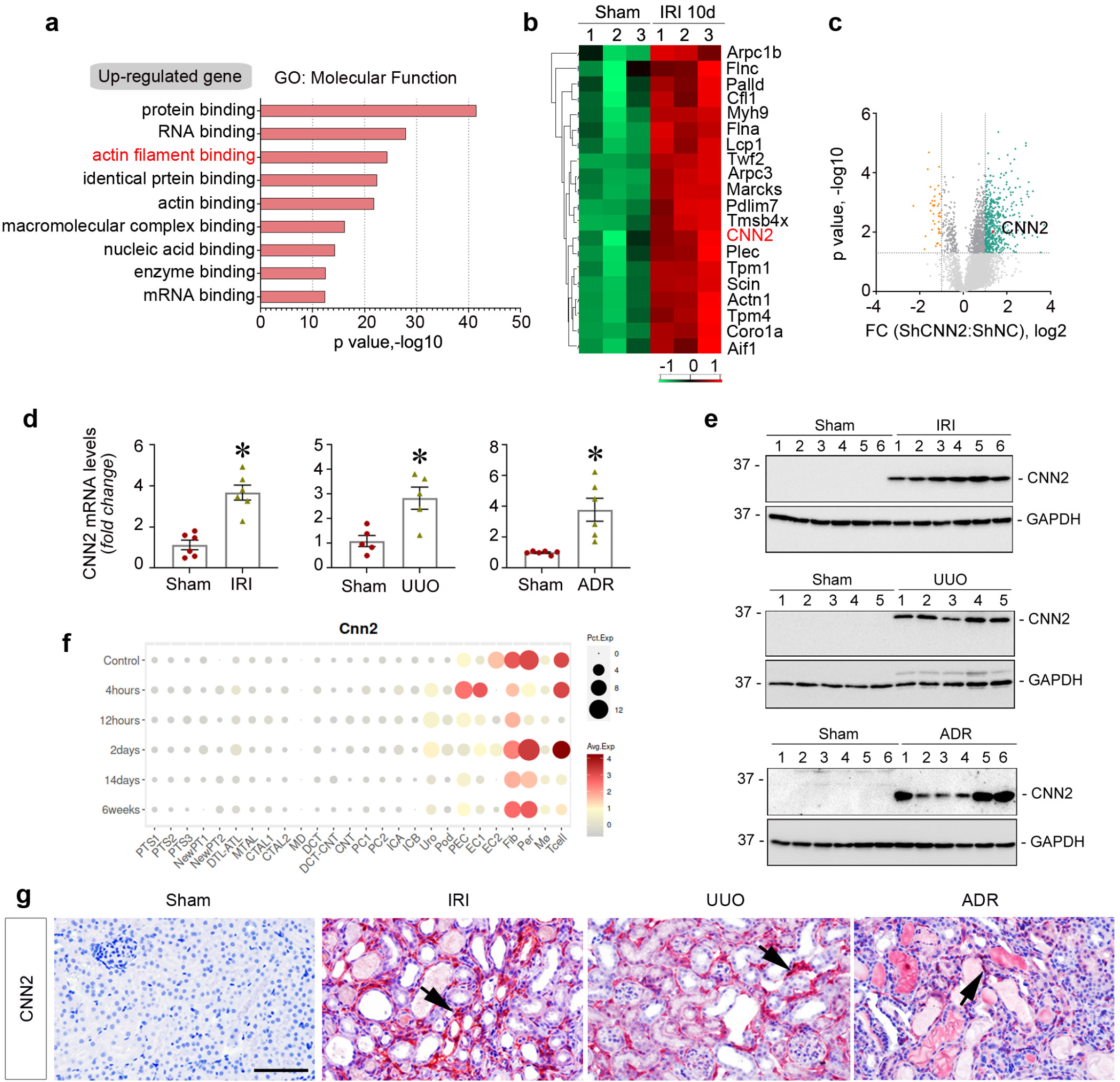
Calponin 2 is a key actin regulatory protein identified in the fibrotic kidneys. **a**) Gene Ontology (GO) enrichment analysis revealed that actin filament binding is listed as one of the top 3 significantly upregulated events under molecular function terms. (**b**) Heatmap of the differentially expressed actin filament regulatory proteins between controls and CKD induced by IRI. (**c**) Volcano plot showed CNN2 is upregulated in the ischemic kidneys after CKD. (**d**) Quantitative RT-PCR analysis revealed that CNN2 mRNA levels were upregulated in the fibrotic kidneys induced by IRI, UUO, and ADR nephropathy, respectively. Graphs are presented as means ± SEM. * *P* < 0.05 (n=5-6). (**e**) Western blot assays demonstrated the expression of CNN2 protein in the fibrotic kidneys induced by IRI, UUO, and ADR nephropathy. Numbers indicate individual animals within each group. (**f**) Single nucleus RNA sequencing showed CNN2 mainly expressed by fibroblasts (Fib) and pericytes (Per) after IRI. (Data were extracted from the online database provided by Dr. Benjamin Humphrey’s laboratory at the Washington University in St. Louis, http://humphreyslab.com/SingleCell/displaycharts.php). (**g**) Immunohistochemical staining showed CNN2 distributions in the fibrotic kidneys after IRI, UUO, and ADR nephropathy, respectively. Arrows indicate positive staining. Scale bar, 50 µm. IRI, ischemia reperfusion injury; UUO, unilateral ureteral obstruction; ADR, Adriamycin.

To establish the clinical relevance of CNN2 expression in human CKD, we immunohistochemically stained kidney biopsy specimens from CKD patients with different etiologies for CNN2. CNN2 was induced in the interstitium of all CKD patients, compared with non-tumor HA controls (Figure 3, a-b). To determine whether CNN2 is detectable in circulation, we analyzed serum CNN2 levels from 75 HA and 84 non-diabetic CKD patients and found the level of serum CNN2 was increased in CKD patients (Figure 3c). Interestingly, at the glomerular filtration rate (GFR) range from 60 to 89ml/min/1.73m^2^, serum CNN2 contents are correlated with low-density lipoprotein-cholesterol (LDL-C) levels (Figure 3d).

**Figure 3.**
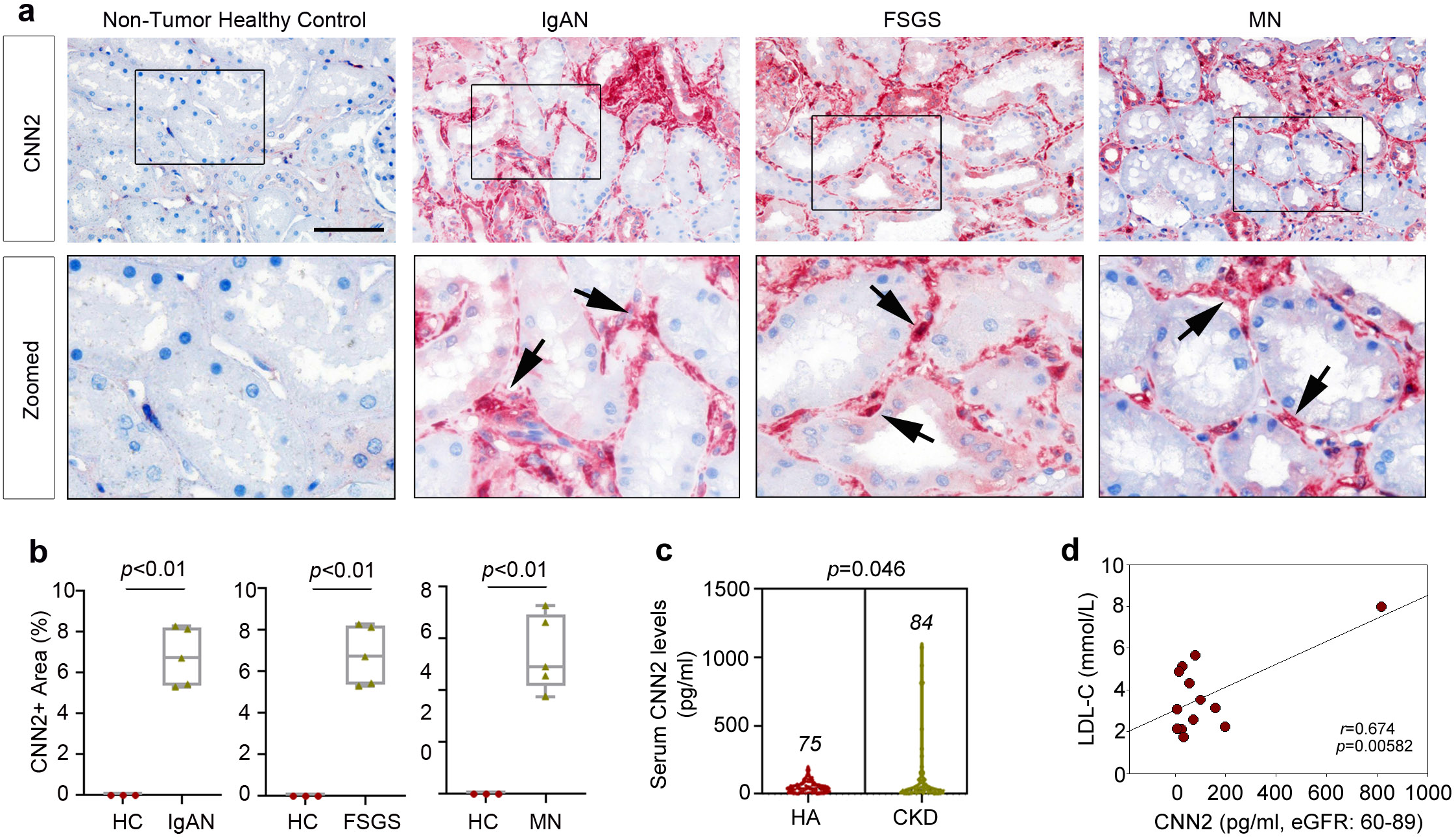
CNN2 expression in CKD patients. (**a, b**) Representative immunohistochemical staining images showed CNN2 expression in the non-tumor normal human kidneys and the kidney biopsy specimens from CKD patients diagnosed with IgA nephropathy (IgAN), focal segmental glomerulosclerosis (FSGS), and membrane nephropathy (MN). Boxed areas are zoomed. Arrows indicated positive staining. Scale bar, 50 μm. Quantitative data are presented (**b**). Graphs are presented as means ± SEM. * *P* < 0.05 (n=5). (**c**) Serum CNN2 levels in healthy adults (n=75) and non-diabetic CKD patients (n=84). Graphs are presented as means ± SEM. (**d**) Correlation between CNN2 and low-density lipoprotein-cholesterol (LDL-C) in CKD patients (GFR, 60-89 ml/min/1.73m^2^).

### CNN2 knockdown alleviates kidney fibrosis

To explore the role of CNN2 in kidney fibrosis, we employed a hydrodynamic gene delivery technique to knock down CNN2 *in vivo* (Figure 4a). Compared to vehicles, qPCR and western blot confirmed that CNN2 induction was reduced in the fibrotic kidneys of ShCNN2 mice (Figure 4b-4c; Supplementary Figure S3a). Immunohistochemical staining consistently showed decreased CNN2 in the interstitium of the fibrotic kidneys (Figure 4d). Then, we examined whether CNN2 knockdown could preserve kidney function and alleviate kidney fibrosis. After IRI, BUN and Scr levels were reduced (Figure 4e-4f), and expression of fibrosis-related genes, including fibronectin (FN), α1 type I collagen (Col1α1), and α1 type III collagen (Col3α1), were reduced in ShCNN2 kidneys compared to vehicles (Figure 4g). Western blots demonstrated reductions of Col1α1, FN, α-smooth muscle actin (α-SMA), and platelet-derived growth factor receptor β (PDGFR-β) in ShCNN2 kidneys (Figures 4h; Supplementary Figure S3b). Immunohistochemical staining illustrated that CNN2 knockdown repressed α-SMA+ myofibroblast activation, Col1α1 deposition, and FN inductions (Figure 4i; Supplementary Figure S3c). Masson trichrome staining (MTS) showed reduced ECM deposition in ShCNN2 kidneys. Additionally, qPCR analysis revealed decreased mRNA levels of secreted cyto-chemokines including monocyte chemoattractant protein-1 (MCP-1), interleukin 6 (IL-6), and tumor necrosis factor-α (TNF-α) in ShCNN2 kidneys (Supplementary Figure S3d). Immunostaining showed fewer CD45+ monocytes and CD3+ T cells in ShCNN2 kidneys (Supplementary Figure S3, e-f). Consistent with the IRI model, CNN2 knockdown exhibited similar effects on reducing ECM deposition and inflammation in the UUO model (Supplementary Figure S4).

**Figure 4:**
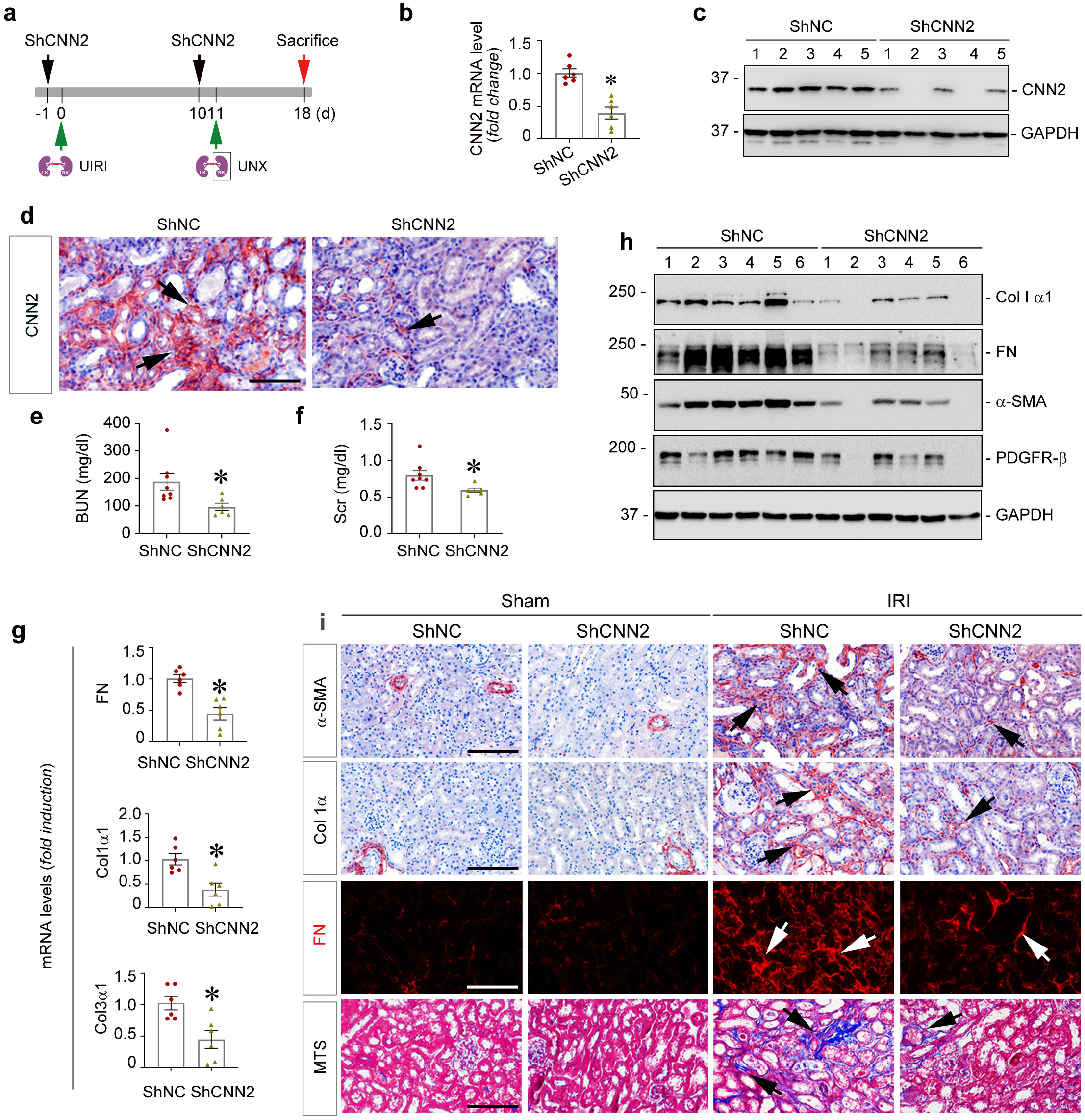
Knockdown of CNN2 alleviates kidney fibrosis. (**a**) Experiment design. The first dose of ShCNN2 plasmid was administrated in mice 1 day (d) before unilateral IRI (UIRI) surgery. At d10 after IRI (1d before unilateral nephrectomy, UNX), the second dose of ShCNN2 plasmid was injected. The mice were sacrificed on d18. (**b**) Quantitative RT-PCR (qPCR) analysis showed the changes of *CNN2* mRNA levels in kidneys of ShNC and ShCNN2 mice after IRI. Graphs are presented as means ± SEM. \**P* < 0.05 (n=6). (**c**) Western blot assay demonstrated CNN2 protein expression in kidneys of ShNC and ShCNN2 mice after IRI. Numbers indicate individual animals within each group. (**d**) Immunohistochemical staining showed CNN2 expression and distribution in kidneys of shNC and shCNN2 mice after IRI. Scale bar, 50 µm. (**e, f**) Blood urea nitrogen (BUN) and serum creatinine (Scr) levels in shCNN2 and shNC mice after IRI. Graphs are presented as means ± SEM. * *P* < 0.05 (n=6-8). (**g**) qPCR analyses revealed the mRNA abundance of *FN, α1 type I collagen (Col1α1)*, and *α1 type III collagen (Col3α1)* in kidneys of shNC and shCNN2 mice after IRI. Graphs are presented as means ± SEM. * *P* < 0.05 (n=6). (**h**) Western blot assay demonstrated Col1α1, FN, α-smooth muscle actin (α-SMA), and platelet-derived growth factor receptor β (PDGFRβ) protein expression in kidneys of shNC and shCNN2 mice after IRI. Numbers indicate individual animals within each group. Graphs are presented as means ± SEM. * *P* <0.05 (n=6). (**i**) Representative micrographs for α-SMA, Col1α1, FN staining, and Masson Trichrome Staining in fibrotic kidneys of shNC and shCNN2 mice after IRI-induced CKD. Arrows indicate positive staining. Scale bar, 50μm.

### Global proteomics reveals FAO is a key pathway after CNN2 knockdown

To better understand the molecular mechanisms of how CNN2 knockdown alleviates CKD, we employed a label-free quantitative approach to profile the proteome landscape of ShNC and ShCNN2 kidneys (Figure 5a). PCA classified ShNC and ShCNN2 kidneys according to their genotype (Figure 5b). A t-test identified 2419 differentially expressed proteins (Permutation FDR 0.05) between ShNC and ShCNN2 mice. Compared to ShNC kidneys, 906 and 1513 proteins were up- and down-regulated in ShCNN2 kidneys (Figure 5, c-d; Supplementary Table S6). The proteomic measurements were reproducible as the protein intensity distribution was similar and the Pearson correlation of the six biological replicates of each group was 0.95 or higher (Supplementary Figure S5, a-b). GO and KEGG enrichment analysis revealed that the metabolic pathway is significantly upregulated in ShCNN2 fibrotic kidneys and FA metabolic process is the most changed event (Figure 5, e and g). KEGG also identified the downregulated pathways and biological processes (Supplementary Figure S5c and S5d). We further investigated the dysregulated metabolic pathways. Among the FA, carbohydrate, and amino acid metabolisms, peroxisome, propanoate, and branched-chain amino acid metabolisms were significantly enriched (Figure 5f). Impressively, about 40 components in the FAO pathway, including Acsm, Acox, and CPT, were simultaneously upregulated in ShCNN2 kidneys compared to ShNC (Figure 5h).

**Figure 5.**
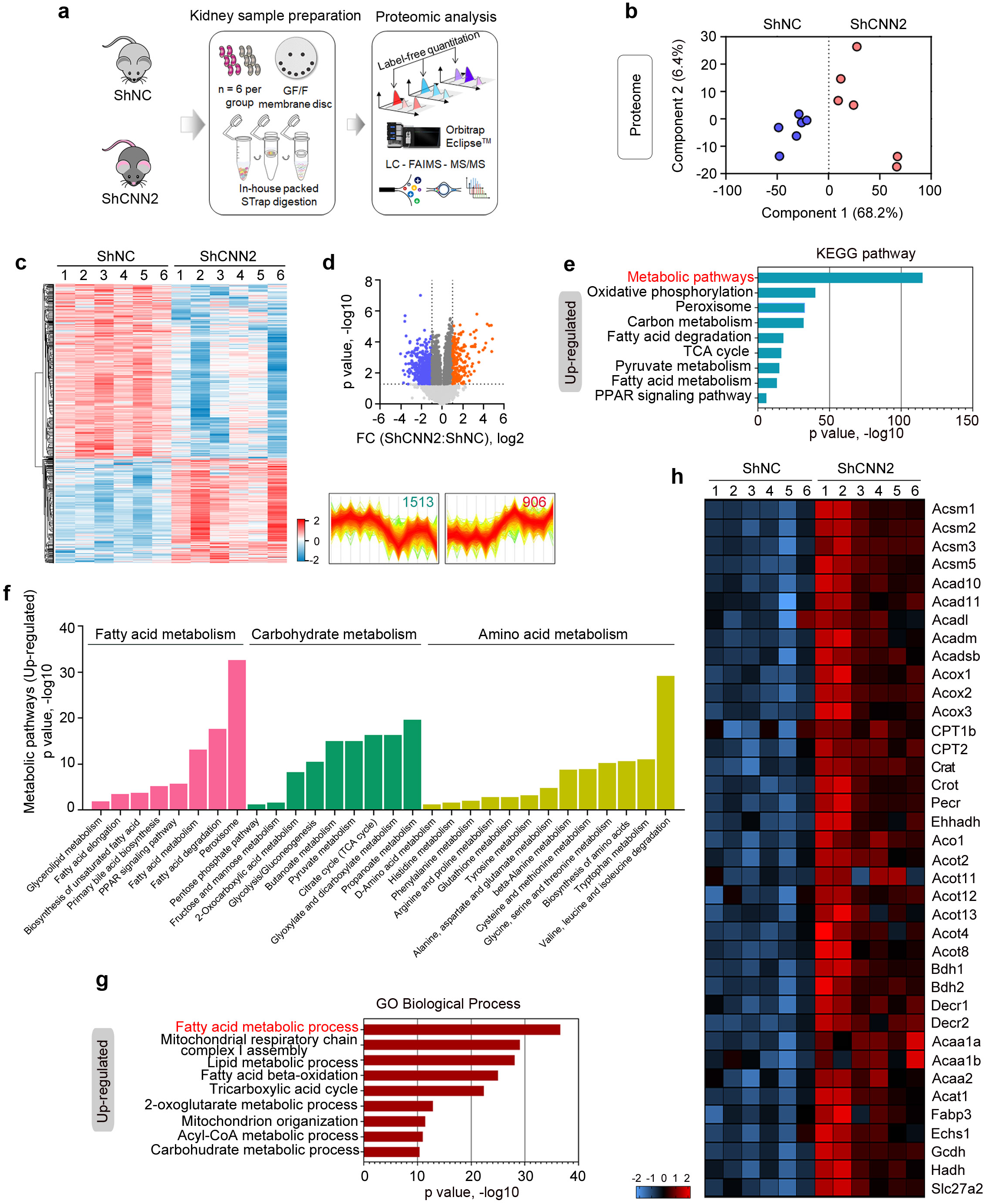
Global proteomics reveals fatty acid oxidation is a key pathway in mediating kidney fibrosis after knockdown of CNN2. (**a**) Experimental workflow of the global proteomic analysis. In each group, kidney samples from six mice were used for mass spectrometry. (**b**) Principal component analysis of global proteomes from ShNC and ShCNN2 fibrotic kidneys after IRI. (**c**) Heatmap of t-test significant proteins (Permutation FDR 0.05). Label-free quantitation (LFQ) intensity of represented proteins were z-scored and plotted according to the color bar. Two clusters of proteins with different patterns of abundance profiles are highlighted in the heatmap. (**d**) Volcano plot showed the differential proteins (Permutation FDR 0.05) between ShNC and ShCNN2 fibrotic kidneys. Up- and down-regulated proteins (fold-change, FC) are colored in red and blue, respectively. (**e, f**) Kyoto Encyclopedia of Genes and Genomes (KEGG) pathway enrichment analysis highlighted upregulated metabolic pathways in fibrotic kidneys from ShNC and ShCNN2 mice after IRI. (**g**) Gene Ontology (GO) biological process terms in each cluster of proteins are plotted with their names and significance. The fatty acid metabolic process is boxed to indicate the group with the largest difference in upregulated proteins. (**h**) Heatmap of the differentially expressed proteins in the fatty acid oxidation pathway in ShNC and ShCNN2 mice fibrotic kidneys after IRI.

### CNN2 modifies lipid accumulation and FAO pathway activity in fibrotic kidneys

We then validated FAO regulation in ShNC and ShCNN2 kidneys. First, we examined the extent of lipid accumulation in fibrotic kidneys. Triglyceride content in sera or kidneys was decreased in ShCNN2 mice compared to ShNC after IRI (Figure 6, a-b). Oil red O staining consistently showed less triglyceride in ShCNN2 kidney tubules (Figure 6c). To enhance the evidence of lipid accumulation in the fibrotic kidneys, we employed an additional marker, perilipin 2 (Plin2), as it is a marker for lipid droplets.^35^ Western blots demonstrated that Plin2 is dramatically decreased in the ShCNN2 kidneys compared to ShNC (Figure 6d; Supplementary Figure S6a). Immunohistochemical staining revealed that Plin2 was predominantly distributed in ShNC kidney tubules (Figure 6e). Because FAO is the key contributor to intracellular ATP levels in tubular cells, we detected ATP consumption in the fibrotic kidneys. ELISA showed the ATP levels were higher in ShCNN2 kidneys than ShNC (Figure 6f).

**Figure 6:**
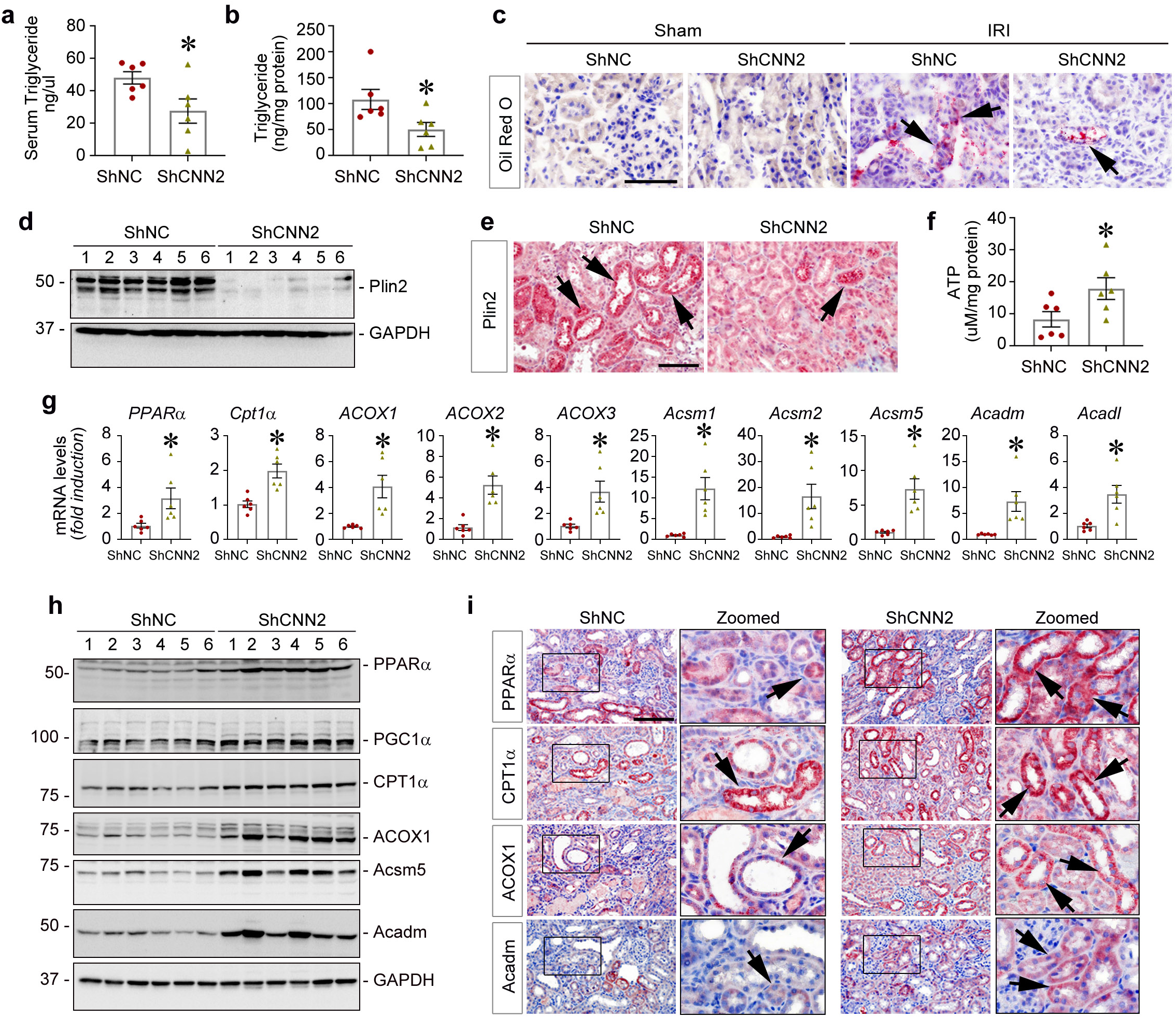
CNN2 modifies lipid accumulation and fatty acid oxidation pathway amid kidney fibrosis. (**a, b**) The enzyme-linked immunosorbent assay (ELISA) showed triglyceride contents in serum (**a**) and kidney (**b**) collected from ShNC and ShCNN2 mice after IRI. Graphs are presented as means ± SEM. * *P* < 0.05 (n=6). (**c**) Representative micrographs of Oil Red-O staining in fibrotic kidneys from ShNC and ShCNN2 mice after IRI. Arrows indicate lipid accumulation in tubules. Scale bar, 50 μm. (**d**) Western blot assay demonstrated perilipin 2 (Plin2) protein expression in ShNC and ShCNN2 mice fibrotic kidneys after IRI. Numbers indicate individual animals within each group. (**e**) Immunohistochemical staining showed Plin2 distributions in fibrotic kidneys from ShNC and ShCNN2 mice after IRI. Arrows indicate positive staining. Scale bar, 50 μm. (**f**) ELISA showed ATP levels in the total kidneys collected from ShNC and ShCNN2 mice after IRI. Graphs are presented as means ± SEM. * *P* < 0.05 (n=6). (**g**) Quantitative RT-PCR analyses revealed the mRNA abundance of *PPARα, CPT1α, ACOX1-3, Acsm1, Acsm2, Acsm5, Acadm*, and *Acadl* in fibrotic kidneys from ShNC and ShCNN2 mice after IRI. Graphs are presented as means ± SEM. * *P* < 0.05 (n=6). (**h**) Western blot assays demonstrated PPARα, PGC1α, CPT1α, ACOX1, Acsm5, and Acadm protein expression in ShNC and ShCNN2 mice fibrotic kidneys after IRI. Numbers indicate individual animals within each group. (**i**) Representative micrographs of PPARα, CPT1α, ACOX1, and Acadm staining in fibrotic kidneys form ShNC and ShCNN2 mice after IRI. Boxed areas are enlarged. Arrows indicate positive staining. Scale bar, 50 μm.

Next, we examined expression of key elements in the FAO pathway in ShNC and ShCNN2 kidneys. qPCR analyses showed that a key transcriptional regulator of FAO, PPARα, was upregulated in ShCNN2 kidneys compared to ShNC (Figure 6g). Meanwhile, the key rate-limiting enzymes of FAO, including CPT1α, ACOX1-3, Acsm1, Acsm2, Acsm5, Acadm, and Acadl were also increased in ShCNN2 kidneys. Western blots confirmed upregulation of PPARα and PPARγ coactivator-1α (PGC-1α) in ShCNN2 kidneys. The downstream targets, including CPT1α, ACOX1, Acsm5, and Acadm were accordingly increased (Figure 6h; Supplementary Figure S6b). Immunohistochemical staining showed upregulation of PPARα, CPT1α, ACOX1, and Acadm in ShCNN2 kidney tubules (Figure 6i). Additionally, CNN2 knockdown also reduced lipid accumulation and FAO defect in the UUO model (Supplementary Figure S7).

### CNN2 enhances CPT1α to preserve kidney function after CKD

To further elucidate how FAO impacts CKD, we applied etomoxir to specifically inhibit CPT1α, the key rate-limiting enzyme in the FAO pathway (Figure 7a). At mRNA and protein levels, etomoxir repressed CPT-1α expression, but ShCNN2 restored its induction (Figure 7, b-c; Supplementary Figure S8a). As predicted, etomoxir increased BUN and Scr levels after IRI. However, under etomoxir treatment, CNN2 knockdown preserved kidney function (Figure 7, d-e). Oil Red O and immunohistochemical staining for Plin2 showed reduced lipid droplet accumulation in ShCNN2 kidney tubules compared to ShNC after etomoxir treatment (Figure 7f). Furthermore, we assessed the extent of kidney fibrosis. qPCR analysis revealed etomoxir increased the mRNA levels of FN, Col1α1, and Col3α1 after IRI, but CNN2 knockdown repressed their induction (Figure 7g). Western blots demonstrated that etoxomir induced FN and α-SMA but ShCNN2 suppressed them (Figure 7h; Supplementary Figure S8b). Immunostaining and MTS showed consistent results on myofibroblast activation and ECM deposition (Figure 7i; Supplementary Figure 8c). Additionally, CNN2 knockdown reduced cyto-chemokine secretion, including MCP-1, IL-6, and TNF-α, and infiltration of CD45+ monocytes after etomoxir treatment (Supplementary Figure S8, d-f).

**Figure 7.**
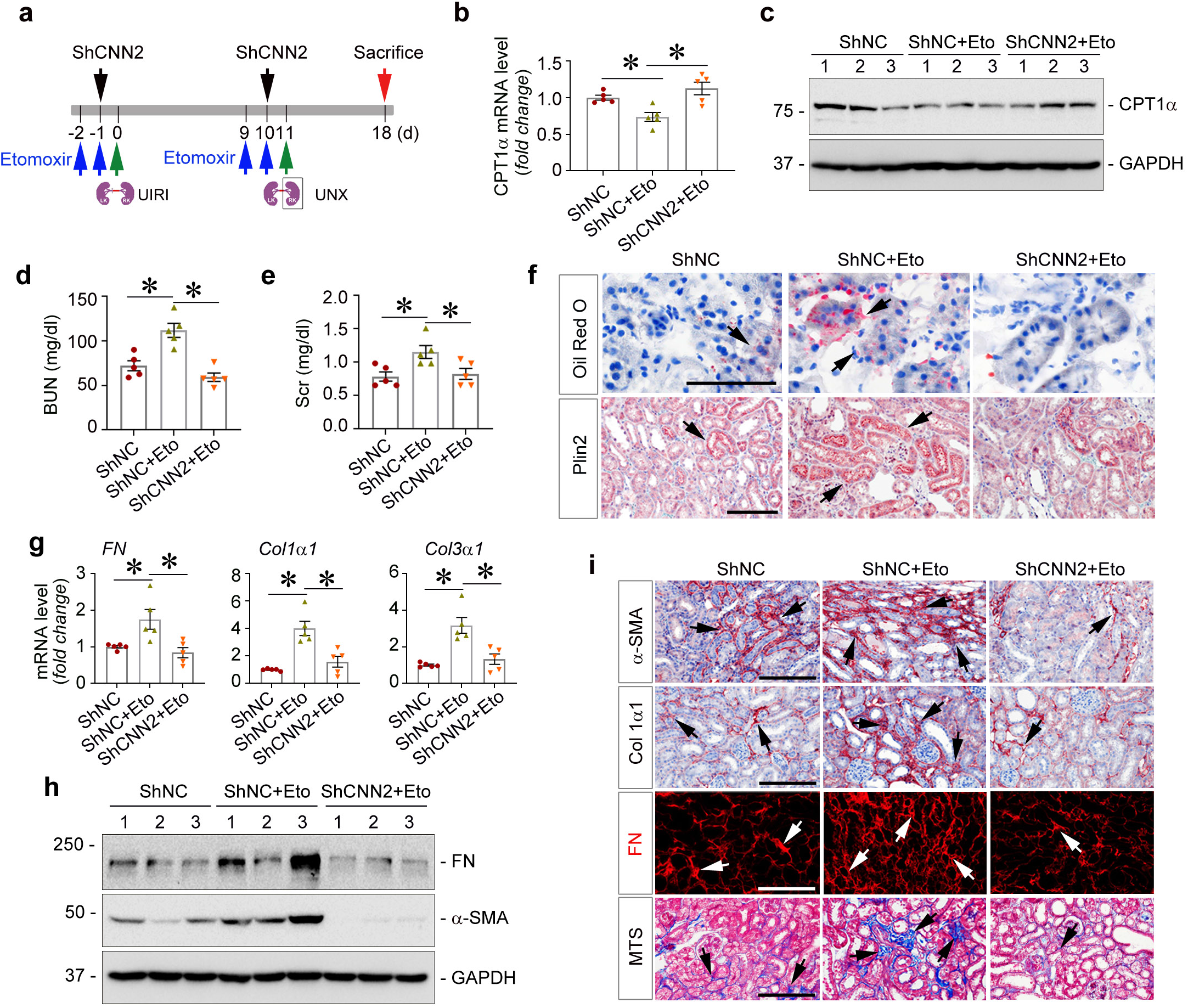
Knockdown of CNN2 enhances CPT1α activity to alleviate kidney fibrosis. (**a**) Experiment design. CPT1α inhibitor, Etomoxir (Eto), was administrated in mice 2 days before unilateral IRI (UIRI) and nephrectomy (UNX), respectively. ShNC and ShCNN2 plasmids were injected twice at day (d) -1 and 10 through the tail vein. The mice were sacrificed on d18. (**b**) Quantitative RT-PCR (qPCR) analysis showed the changes of *CPT1α* mRNA levels in the fibrotic kidneys of ShNC mice, ShNC mice that received Etomoxir, and ShCNN2 mice that received Etomoxir after IRI. Graphs are presented as means ± SEM. * *P* <0.05 (n=5). (**c**) Western blot assay demonstrated CPT1α protein expression in the fibrotic kidneys of ShNC mice, ShNC mice that received Etomoxir, and ShCNN2 mice that received Etomoxir after IRI. Numbers indicate individual animals within each group. (**d, e**) Blood urea nitrogen (BUN) and serum creatinine (Scr) levels in ShNC mice, ShNC mice that received Etomoxir, and ShCNN2 that received Etomoxir after IRI. Graphs are presented as means ± SEM. * *P* < 0.05 (n=5). (**f**) Representative micrographs for Oil Red O Staining and Plin2 immunohistochemical staining showed lipid accumulations in fibrotic kidneys of ShNC mice, ShNC mice that received Etomoxir, and ShCNN2 mice that received Etomoxir after IRI. Arrows indicate positive staining. Scale bar, 50 µm. (**g**) qPCR analyses revealed the mRNA abundance of *FN, α1 type I collagen*, and *α1 type III collagen* in fibrotic kidneys of ShNC mice, ShNC mice that received Etomoxir, and ShCNN2 mice that received Etomoxir after IRI. Graphs are presented as means ± SEM. * *P* < 0.05 (n=5). (**h**) Western blot assay demonstrated FN and α-SMA proteins expression in fibrotic kidneys of ShNC mice, ShNC mice that received Etomoxir, and ShCNN2 mice that received Etomoxir after IRI. Numbers indicate individual animals within each group. (**i**) Representative micrographs for immunostaining of α-SMA, Col1α1, fibronectin (FN), and Masson Trichrome Staining (MTS) showed myofibroblast activation and collagen deposition in fibrotic kidneys of ShNC mice, ShNC mice that received Etomoxir, and ShCNN2 mice that received Etomoxir after IRI. Arrows indicate positive staining. Scale bar, 50 µm.

**Figure 8.**
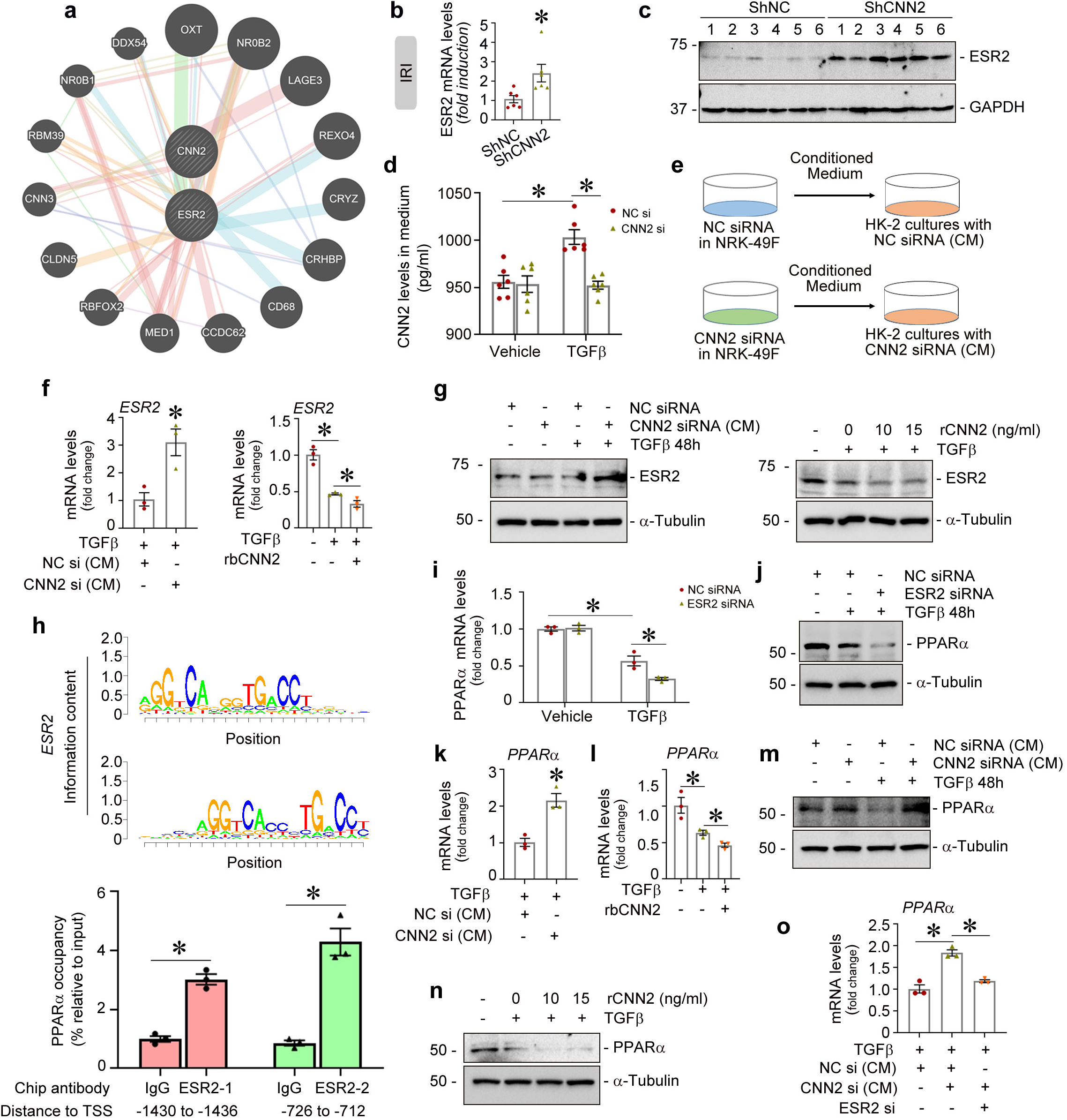
Knockdown of CNN2 promotes ESR2 binding PPARα to enhance fatty acid oxidation. (**a**) Based on the GeneMANIA database, the human protein-protein interaction analysis showed CNN2 interacts with estrogen receptor 2 (ESR2). (**b)** Quantitative RT-PCR (qPCR) analysis showed the abundance of *ESR2* mRNA levels in ShNC and ShCNN2 mice fibrotic kidneys after IRI. Graphs are presented as means ± SEM. * *P* <0.05 (n=6). (**c**) Western blot assay demonstrated ESR2 protein expression in fibrotic kidneys of ShNC and ShCNN2 mice after IRI. Numbers indicate individual animals within each group. (**d**) ELISA showed CNN2 levels in the conditioned medium (CM) collected from the cultured fibroblasts after knockdown of CNN2 under TGF-β stress. Graphs are presented as means ± SEM. * *P* <0.05 (n=6). (**e**) Schematic diagram showed CNN2-deprived CM from the cultured fibroblasts or recombinant CNN2 protein were employed to treat human kidney proximal tubular cells (HK-2). (**f)** qPCR showed *ESR2* expression in HK-2 cells after incubation with CNN2-deprived CM or human recombinant CNN2 protein. Graphs are presented as means ± SEM. * *P* <0.05 (n=3). (**g**) Western blot assay demonstrated ESR2 expression in HK-2 cells after incubation with CNN2-deprived CM or human recombinant CNN2 protein. (**h**) ChIP-qPCR assay showed ESR2 binds to putative sequences in the promoter of *PPARα* genes. HK-2 cells were treated with ESR2 agonist (LY500307) for 3 hours. Chromatin preparations from the cells were immunoprecipitated using an anti-ESR2 antibody, and co-precipitated DNA fragments were amplified using primers specific for the PPARα promoter. Graphs are presented as means ± SEM. * *P* <0.05 (n=3). The upper panel showed representative DNA sequence logo representing the binding motif of the ESR2 gene. (**i**) qPCR analyses revealed the mRNA abundances of *PPARα* were reduced in HK-2 cells after knockdown of ESR2 under TGFβ1 stress, compared with scramble controls. Graphs are presented as means ± SEM. * *P* < 0.05 (n=3). (**j**) Western blot assay demonstrated that knockdown of ESR2 repressed PPARα expression in HK-2 cells under TGFβ1 stress, compared with scramble controls. (**k, l**) qPCR analyses revealed the mRNA abundances of *PPARα* were increased in HK-2 cells by CNN2-deprived CM (**k**) but were repressed by CNN2 recombinant protein (**l**) under TGFβ1 stress, compared with controls. Graphs are presented as means ± SEM. * *P* < 0.05 (n=3). (**m, n**) Western blot assays demonstrated that CNN2-deprived CM enhanced PPARα expression (**m**) but CNN2 recombinant protein repressed PPARα inductions (**n**) in HK-2 cells under TGFβ1 stress, compared with controls. (**o**) qPCR analyses revealed the mRNA abundances of *PPARα* were induced by CNN2-deprived CM but were repressed after knockdown of ESR2 under TGFβ1 stress. Graphs are presented as means ± SEM. * *P* <0.05 (n=3).

### CNN2 knockdown promotes ESR2 binding to PPARα to enhance transcription of FAO genes

Given that CNN2 knockdown affected multiple FAO genes and its regulator PPARα expression (Figure 6), we hypothesized that there is a common mechanism that controls their activation, so we performed bioinformatics analyses. Based on GeneMANIA database, the human protein-protein network analysis indicated that estrogen receptor 2 (ESR2) is a strong interactor of CNN2 (Figure 8a). qPCR analyses showed that ESR2 mRNA were increased in ShCNN2 kidneys compared to ShNC after IRI and UUO (Figure 8b; Supplementary Figure S9a). Western blots demonstrated increased ESR2 induction in ShCNN2 kidneys (Figure 8c; Supplementary Figure S9b). Under TGF-β stress *in vitro*, ELISA showed that CNN2 levels in fibroblast (NRK-49F) conditioned medium (CM) were reduced after CNN2 knockdown (Figure 8d; Supplementary Figure S9c). We then employed CNN2-deprived CM and CNN2 recombinant protein to treat human kidney proximal tubular cells (HK-2) (Figure 8e). qPCR analysis revealed that ESR2 mRNA was induced by CNN2-deprived CM but was repressed by CNN2 recombinant protein after TGF-β stimulation (Figure 8f). Western blot assays confirmed ESR2 expression at the protein level (Figure 8g). Because ESR2 is a member of the superfamily of nuclear receptor transcription factors, to examine whether ESR2 directly regulates PPARα, we treated HK-2 cells with an ESR2 agonist (LY500307) and then performed a chromatin immunoprecipitation (ChIP) assay using an anti-ESR2 antibody. Through the JASPAR database, we predicted there are at least 12 putative binding sites for ESR2 on the PPARα gene promoter region (Supplementary Figure S9d). The co-precipitated chromatin DNA fragments extracted from the cells were amplified by qPCR using two pairs of primers for the PPARα promoter region. We found that the ESR2 agonist promoted ESR2 binding to the putative sites on the PPARα gene promoter (Figure 8h). *In vitro*, ESR2 knockdown in HK-2 cells repressed PPARα at both mRNA and protein levels under TGF-β stress (Figure 8, i-j; Supplementary Figure S9e). Interestingly, PPARα levels were increased after incubation with CNN2-deprived CM in HK-2 cells under TGF-β stress (Figure 8, k and m). Conversely, CNN2 recombinant protein repressed PPARα expression (Figure 8, l and n). However, ESR2 knockdown abolished PPARα induction upon incubation with CNN2-deprived CM (Figure 8o).

### CNN2 knockdown in fibroblasts promotes FAO in tubular cells

To study how CNN2 regulates FAO in tubular cells, we incubated HK-2 with CNN2-deprived CM. qPCR analysis revealed that the key rate-limiting enzymes including CPT1α, ACOX1, Acsm1, and Acsm5 mRNA levels were induced by CNN2-deprived CM under TGF-β stress (Figure 9a). Consequently, CNN2-deprived CM reduced FN induction and inhibited α-SMA expression (Figure 9b). However, ESR2 knockdown largely abolished induction of CPT1α, ACOX1, and Acsm1 and increased FN and α-SMA (Figure 9, c-d). To confirm that CNN2 mediates kidney fibrosis by affecting the FAO pathway in tubules, HK-2 cells were incubated with CNN2-deprived CM followed by etomoxir under TGF-β stress. Western blot assays demonstrated that etomoxir abolished the beneficial effects of CNN2-deprived CM on reducing induction of FN and α-SMA (Figure 9e). In comparison, HK-2 cells treated with human recombinant CNN2 protein repressed mRNA levels of PGC1α, CPT1α, ACOX1, and Acsm5 after TGF-β stimulation (Figure 9f). Similarly, CNN2 recombinant protein increased FN and α-SMA induction (Figure 9g). In a separate experiment, HK-2 cells were treated with CNN2 recombinant protein followed by the PPARα agonist, fenofibrate. Under TGF-β stress, western blots demonstrated that fenofibrate ameliorated CNN2-induced FN and α-SMA (Figure 9h).

**Figure 9:**
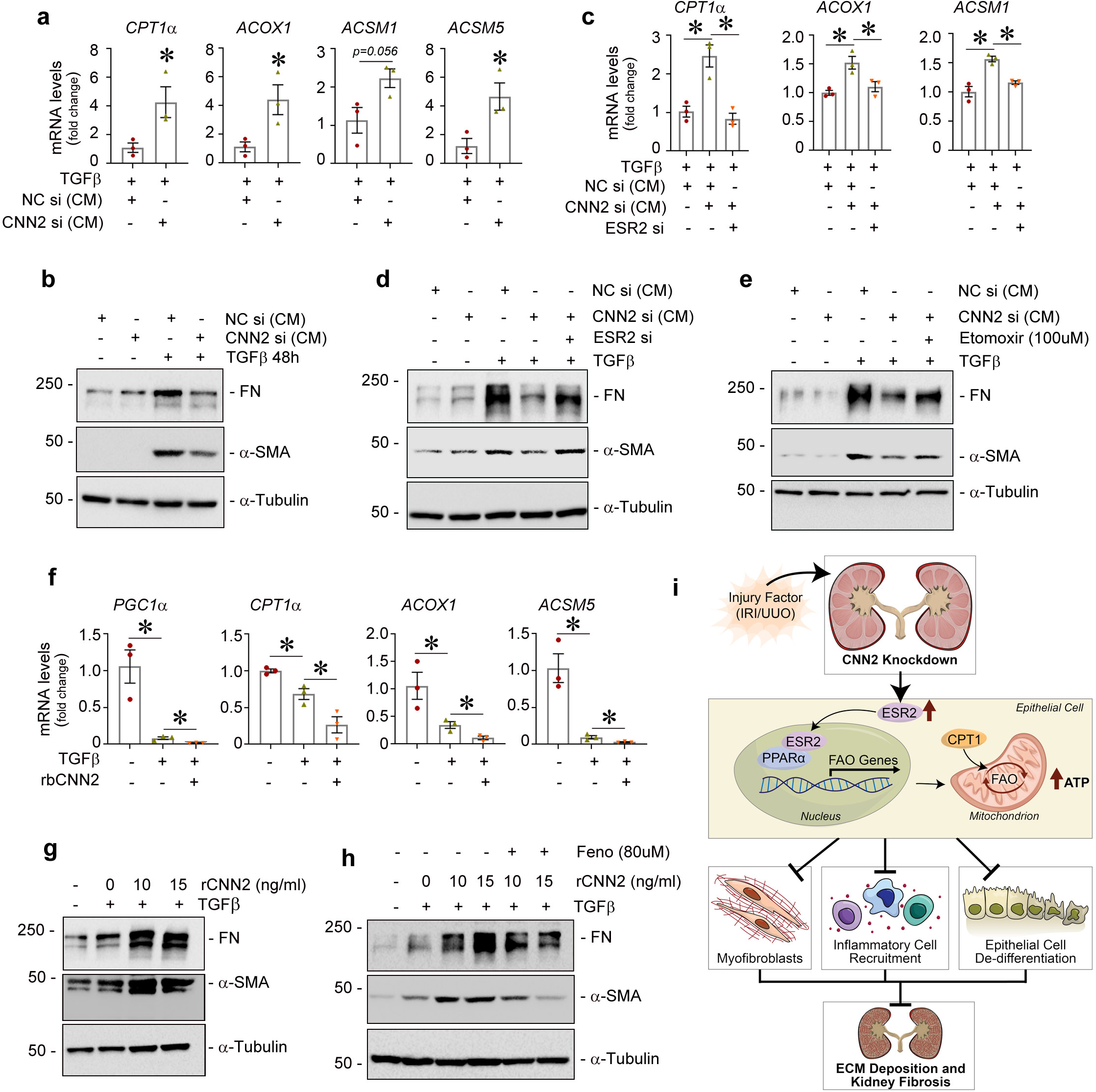
Knockdown of CNN2 alleviates defective fatty acid oxidation and ECM deposition in tubular cells. (**a**) qPCR analyses revealed the mRNA abundances of *CPT1α, ACOX1, Acsm1*, and *Acsm5* were increased in HK-2 cells incubated with CNN2-deprived CM under TGFβ1 stress, compared with controls. Graphs are presented as means ± SEM. * *P* < 0.05 (n=3). (**b**) Western blot assay demonstrated that CNN2-deprived CM reduced the inductions of FN and α-SMA in HK-2 cells under TGFβ1 stress, compared with controls. (**c**) qPCR analyses revealed the mRNA abundances of *CPT1α, ACOX1*, and *Acsm1* were increased by CNN2-deprived CM in HK-2 cells but were repressed after knockdown of ESR2 under TGFβ1 stress, compared with controls. Graphs are presented as means ± SEM. * *P* < 0.05 (n=3). (**d**) Western blot assays demonstrated CNN2-deprived CM alleviated FN and α-SMA inductions in HK-2 cells under TGFβ1 stress but they were increased after knockdown of ESR2, compared with controls. (**e**) Under TGFβ1 stress, HK-2 cells were incubated with CNN2-deprived CM and followed by CPT1α inhibitor Etomoxir (100µM). Western blot assays showed Etomoxir induced FN and α-SMA expression after incubation with CNN2-deprived CM. (**f**) qPCR analyses revealed the mRNA abundances of *PGC1α, CPT1α, ACOX1*, and *Acsm5* were reduced in HK-2 cells incubated with CNN2 recombinant protein under TGFβ1 stress. Graphs are presented as means ± SEM. * *P* < 0.05 (n=3). (**g**) Western blot assay demonstrated that CNN2 recombinant induced FN and α-SMA expression in HK-2 cells after TGFβ1 stimulations. (**h**) Under TGFβ1 stress, HK-2 cells were treated with CNN2 recombinant proteins followed by PPARα agonist Fenofibrate (Feno, 80µM). Western blot assays showed Fenofibrate decreased FN and α-SMA expression after treated with CNN2 recombinant protein. (**i**) Schematic diagram depicts knockdown of CNN2 promotes ESR2 binding to PPARα to transcriptionally regulate the FAO genes to halt kidney fibrosis.

## Discussion

This study elucidated CKD pathogenesis from the perspective of cellular mechanics and metabolism interactions (Figure 9i). We demonstrated that CNN2, a prominent actin stabilizer, determines kidney fibrosis through modulating FA metabolism as evident by: 1) CNN2 knockdown alleviated kidney fibrosis in CKD models, 2) if the FAO pathway was activated, then less lipid accumulated in ShCNN2 fibrotic kidneys, 3) CNN2 knockdown preserved kidney functions after inhibiting the key rate-limiting enzyme, CPT-1α, 4) CNN2’s interactor, ESR2, transcriptionally mediates the regulator of the FAO pathway, PPARα, and 5) serum CNN2 is increased and is associated with LDL-C levels in CKD patients. These results provide rationale for expanding the paradigm of mechano-metabolic programming after CKD.

In mice and human fibrotic kidneys, CNN2 is mainly expressed in interstitial fibroblasts or pericytes (Figure 2 and 3). Due to mechanical changes during CKD, injured kidney cells receive information from the microenvironment to coordinate their behavior at local and systemic levels. Our results indicate that CNN2 affects both these levels in CKD. At the local level, it serves as an actin stabilizer. Under physiological conditions, CNN2 controls cell shape, polarity, migration, and establishes intercellular contacts to support organ architecture.^5^ After CKD, fibrotic niche formation is a critical change in kidneys.^30^ The densities, sizes, and cellular/non-cellular components of niches determine the severity of kidney fibrosis. Amid these processes, all kidney cells experience pathological external forces. These highly varied forces may mediate individual cell phenotypes and kidney architecture changes.^4^ Our data show that CNN2 knockdown alleviates kidney fibrosis after ischemic and obstructive CKD (Figure 4; Supplementary Figure S4). From the aspect of mechanical changes, dysregulated CNN2 reduces the injured cells’ ability to quickly reinforce their mechanosensitive structures and cytoskeleton so they do not respond to forces appropriately. After CKD, actin cytoskeleton disarrangement causes aberrant activation of fibroblasts, excessive ECM deposition, and tubular atrophy. At the systemic level, CNN2 is a soluble factor-like protein to mediate mechanical and chemical signal interactions. Rapid CNN2 degradation is required for cytokines to inhibit malignant tumor growth.^36^ A recent study also indicated that CNN2 could be ubiquitylated and removed for efficient lysophagy during lysosomal membrane permeabilization.^37^ CNN2 was increased in CKD patients’ serum, and CNN2 expression is associated with LDL-C levels in stage 2 CKD patients (Figure 3). This phenomenon leaves us questioning, does CNN2 affect lipid or FA metabolism reprogramming in fibrotic kidneys?

Metabolic reprogramming is a hallmark of malignant transformation.^38^ This process includes activating signal transduction cascades, key metabolic enzymes, and transcriptional regulation of genes encoding metabolic enzymes.^39^ Although the interaction between the actin cytoskeleton and metabolic enzymes was observed decades ago,^40^ most studies investigating cellular mechanics remain focused on supporting organ architecture rather than metabolic reprogramming. As a highly metabolically active organ,^41^ kidney metabolism is markedly disturbed after CKD (Figure 1). Our proteomics data suggest that all metabolic pathways, including FA, carbohydrates, and amino acids, and about 40 FAO-related enzymes were upregulated in ShCNN2 kidneys (Figure 5). Specifically, gene expression of the key regulators of FAO (PPARα) and rate-limiting enzymes (Acsm1, 2, 5, CPT1a, ACOX1-3, Acadm, Acadl, and Acad10) were upregulated in ShCNN2 kidney TECs. Although CNN2 is expressed in fibroblasts/pericytes, communication between TECs and fibroblasts is critical in determining CKD severity.^42^ As the largest cell population of kidneys, TECs are not bystanders in kidney fibrosis.^30^ TECs are the main energy consuming cells, and FAO is their chief ATP source in kidneys. Lower FAO in TECs, followed by intracellular lipid accumulation, profoundly switches TECs into a profibrotic phenotype and leads to kidney fibrosis.^43,44^ Indeed, intracellular triglyceride levels were reduced after CNN2 knockdown (Figure 6; Supplementary Figure S7). CNN2 knockdown rescued CPT1α inhibitor-induced kidney fibrosis by stimulating FA import into the mitochondria and initiating mitochondrial fission. Enhanced FAO provides more ATP to maintain the structural integrity of actin filaments.

The most novel finding of this study is that ESR2 is a bridge that connects CNN2 and PPARα to modulate FAO after CKD (Figure 8), which we determined via several lines of evidence. First, ESR2 interacts with CNN2. Second, as a nuclear transcription factor, ESR2 harbors similar conserved sequences with PPARα in its promoter regions, and ESR2 agonist promotes their binding. Third, CNN2 knockdown alleviated kidney fibrosis, whereas knockdown of both CNN2 and ESR2 abolished these protective effects and repressed FAO. Consistent with the protective role of ESR2 in cancer, neuropathies, cardiovascular disease, osteoporosis, menopause, and metabolic disorders,^45^ the finding that CNN2 controls ESR2 to influence FAO and kidney fibrosis provides new insight into how mechanical signals contribute to CKD development; although, FAO is positioned at the crossroad of multiple chemical signals such as TGF-β, Krüppel-like factor 15, and CD36.^46-48^ These studies also provided a theoretical base for applying a biosensor to adjust the dynamic tension of cytoskeletal filaments to treat CKD in the future.

Despite the robust evidence we report to demonstrate how the actin stabilizer CNN2 regulates metabolism, several unanswered questions remain. First, does CNN2 knockdown affect metabolic reprogramming of kidney fibroblasts? Second, does altered glucose metabolism contribute to kidney fibrosis in ShCNN2 kidneys, given that kidneys are important for glucose reabsorption, production, and utilization? Third, as we did not generate CNN2 genetic mouse models, would genetic knockout of CNN2 have the same effects *in vivo*?

In summary, our investigation of cellular mechanics and metabolism interactions identified that the knockdown of actin stabilizer, CNN2, alleviated kidney fibrosis by activating FA metabolism. The new insights on modulating cellular mechanical properties to tune energy production in response to formation of the kidney fibrotic microenvironment supplement our understanding of CKD pathogenesis. The possibility of employing approaches to target mechano-metabolic mechanisms may shed light on treating CKD.

## Data availability

Raw Mass Spectrometry data were deposited in MassIVE with the data set identifier MSV000090189.

## Acknowledgments

This work was supported by the National Institutes of Health (NIH) grants DK116816, DK128529, and DK132059. We are grateful to Dr. Benjamin Humphreys lab at Washington University in St. Louis for using their online database; Dr. Shijia Liu lab at the Affiliated Hospital of Nanjing University of Chinese Medicine for sharing the serum metabolomics data; and Dr. Justin D. Radolf at the University of Connecticut, School of Medicine for reviewing the manuscript.

## Supplementary Material

Supplementary File (PDF)

## Detailed Methods

**Figure S1**: Proteomics and metabolomics profile the landscape of CKD.

**Figure S2**: Calponin 2 is upregulated in the fibrotic kidneys after CKD.

**Figure S3**: Knockdown of CNN2 alleviates kidney fibrosis induced by IRI.

**Figure S4**: Knockdown of CNN2 ameliorates obstructive CKD in mice.

**Figure S5**: Proteomics reveals knockdown of CNN2 activates FAO in the fibrotic kidneys after IRI.

**Figure S6**: Knockdown of CNN2 decreases lipid accumulation and activates FAO pathway after IRI.

**Figure S7**: Knockdown of CNN2 decreases lipid accumulation and activates FAO after UUO.

**Figure S8**: Knockdown of CNN2 enhances CPT1α activity to alleviate kidney fibrosis.

**Figure S9**: Knockdown of CNN2 promotes ESR2 binding to PPARα.

**Table S1**: The demographic and clinical data of the patients (Kidney biopsy).

**Table S2**: The demographic and clinical data of the participants (Metabolomics).

**Table S3**: Nucleotide sequences of the primers used for qRT-PCR.

**Table S4**: The information of the applied primary and secondary antibodies.

**Table S5**: Differentially expressed metabolites in healthy adults and non-diabetic CKD patients,

**Table S6**: Differentially expressed proteins in ShNC and ShCNN2 kidneys after IRI.

## References

1 Foreman, K. J. et al. Forecasting life expectancy, years of life lost, and all-cause and cause-specific mortality for 250 causes of death: reference and alternative scenarios for 2016-40 for 195 countries and territories. Lancet 392, 2052–2090, doi:10.1016/S0140-6736(18)31694-5 (2018).

2 Kalantar-Zadeh, K., Jafar, T. H., Nitsch, D., Neuen, B. L. & Perkovic, V. Chronic kidney disease. Lancet 398, 786–802, doi:10.1016/S0140-6736(21)00519-5 (2021).

3 Li, L., Fu, H. & Liu, Y. The fibrogenic niche in kidney fibrosis: components and mechanisms. Nat Rev Nephrol 18, 545–557, doi:10.1038/s41581-022-00590-z (2022).

4 Wells, R. G. Tissue mechanics and fibrosis. Biochim Biophys Acta 1832, 884–890, doi:10.1016/j.bbadis.2013.02.007 (2013).

5 DeWane, G., Salvi, A. M. & DeMali, K. A. Fueling the cytoskeleton - links between cell metabolism and actin remodeling. J Cell Sci 134, doi:10.1242/jcs.248385 (2021).

6 Liu, R. & Jin, J. P. Calponin isoforms CNN1, CNN2 and CNN3: Regulators for actin cytoskeleton functions in smooth muscle and non-muscle cells. Gene 585, 143–153, doi:10.1016/j.gene.2016.02.040 (2016).

7 Moazzem Hossain, M., Wang, X., Bergan, R. C. & Jin, J. P. Diminished expression of h2-calponin in prostate cancer cells promotes cell proliferation, migration and the dependence of cell adhesion on substrate stiffness. FEBS Open Bio 4, 627–636, doi:10.1016/j.fob.2014.06.003 (2014).

8 Hu, J. et al. Knockdown of calponin 2 suppressed cell growth in gastric cancer cells. Tumour Biol 39, 1010428317706455, doi:10.1177/1010428317706455 (2017).

9 Huang, Q. Q. et al. Deletion of calponin 2 in macrophages attenuates the severity of inflammatory arthritis in mice. Am J Physiol Cell Physiol 311, C673–C685, doi:10.1152/ajpcell.00331.2015 (2016).

10 Plazyo, O., Liu, R., Moazzem Hossain, M. & Jin, J. P. Deletion of calponin 2 attenuates the development of calcific aortic valve disease in ApoE(-/-) mice. J Mol Cell Cardiol 121, 233–241, doi:10.1016/j.yjmcc.2018.07.249 (2018).

11 Hsieh, T. B., Feng, H. Z. & Jin, J. P. Deletion of Calponin 2 Reduces the Formation of Postoperative Peritoneal Adhesions. J Invest Surg 35, 517–524, doi:10.1080/08941939.2021.1880672 (2022).

12 Kang, X. et al. Lentivirus-mediated shRNA Targeting CNN2 Inhibits Hepatocarcinoma in Vitro and in Vivo. Int J Med Sci 15, 69–76, doi:10.7150/ijms.21113 (2018).

13 Evers, T. M. J., Holt, L. J., Alberti, S. & Mashaghi, A. Reciprocal regulation of cellular mechanics and metabolism. Nat Metab 3, 456–468, doi:10.1038/s42255-021-00384-w (2021).

14 Li, Y., Sha, Z. & Peng, H. Metabolic Reprogramming in Kidney Diseases: Evidence and Therapeutic Opportunities. Int J Nephrol 2021, 5497346, doi:10.1155/2021/5497346 (2021).

15 Tran, M. T. et al. PGC1alpha drives NAD biosynthesis linking oxidative metabolism to renal protection. Nature 531, 528–532, doi:10.1038/nature17184 (2016).

16 Bays, J. L., Campbell, H. K., Heidema, C., Sebbagh, M. & DeMali, K. A. Linking E-cadherin mechanotransduction to cell metabolism through force-mediated activation of AMPK. Nat Cell Biol 19, 724–731, doi:10.1038/ncb3537 (2017).

17 Console, L. et al. The Link Between the Mitochondrial Fatty Acid Oxidation Derangement and Kidney Injury. Front Physiol 11, 794, doi:10.3389/fphys.2020.00794 (2020).

18 Miguel, V. et al. Renal tubule Cpt1a overexpression protects from kidney fibrosis by restoring mitochondrial homeostasis. J Clin Invest 131, doi:10.1172/JCI140695 (2021).

19 Kang, H. M. et al. Defective fatty acid oxidation in renal tubular epithelial cells has a key role in kidney fibrosis development. Nat Med 21, 37–46, doi:10.1038/nm.3762 (2015).

20 Han, S. H. et al. PGC-1alpha Protects from Notch-Induced Kidney Fibrosis Development. J Am Soc Nephrol 28, 3312–3322, doi:10.1681/ASN.2017020130 (2017).

21 Khan, S. et al. Kidney Proximal Tubule Lipoapoptosis Is Regulated by Fatty Acid Transporter-2 (FATP2). J Am Soc Nephrol 29, 81–91, doi:10.1681/ASN.2017030314 (2018).

22 Baek, J., He, C., Afshinnia, F., Michailidis, G. & Pennathur, S. Lipidomic approaches to dissect dysregulated lipid metabolism in kidney disease. Nat Rev Nephrol 18, 38–55, doi:10.1038/s41581-021-00488-2 (2022).

23 Iwai, N. et al. Association between SAH, an acyl-CoA synthetase gene, and hypertriglyceridemia, obesity, and hypertension. Circulation 105, 41–47, doi:10.1161/hc0102.101780 (2002).

24 Szeto, H. H. Pharmacologic Approaches to Improve Mitochondrial Function in AKI and CKD. J Am Soc Nephrol 28, 2856–2865, doi:10.1681/ASN.2017030247 (2017).

25 He, M. et al. Identification and characterization of new long chain acyl-CoA dehydrogenases. Mol Genet Metab 102, 418–429, doi:10.1016/j.ymgme.2010.12.005 (2011).

26 Jang, H. S., Noh, M. R., Kim, J. & Padanilam, B. J. Defective Mitochondrial Fatty Acid Oxidation and Lipotoxicity in Kidney Diseases. Front Med (Lausanne) 7, 65, doi:10.3389/fmed.2020.00065 (2020).

27 Chau, B. N. et al. MicroRNA-21 promotes fibrosis of the kidney by silencing metabolic pathways. Sci Transl Med 4, 121ra118, doi:10.1126/scitranslmed.3003205 (2012).

28 Zhou, D. & Liu, Y. Renal fibrosis in 2015: Understanding the mechanisms of kidney fibrosis. Nat Rev Nephrol 12, 68–70, doi:10.1038/nrneph.2015.215 (2016).

29 Percie du Sert, N. et al. The ARRIVE guidelines 2.0: Updated guidelines for reporting animal research. PLoS Biol 18, e3000410, doi:10.1371/journal.pbio.3000410 (2020).

30 Fu, H. et al. The hepatocyte growth factor/c-met pathway is a key determinant of the fibrotic kidney local microenvironment. iScience 24, 103112, doi:10.1016/j.isci.2021.103112 (2021).

31 Zhou, D. et al. Non-canonical Wnt/calcium signaling is protective against podocyte injury and glomerulosclerosis. Kidney Int 102, 96–107, doi:10.1016/j.kint.2022.02.029 (2022).

32 Lin, Y. H. et al. Global Proteome and Phosphoproteome Characterization of Sepsis-induced Kidney Injury. Mol Cell Proteomics 19, 2030–2047, doi:10.1074/mcp.RA120.002235 (2020).

33 Liu, S. et al. Serum integrative omics reveals the landscape of human diabetic kidney disease. Mol Metab 54, 101367, doi:10.1016/j.molmet.2021.101367 (2021).

34 Kirita, Y., Wu, H., Uchimura, K., Wilson, P. C. & Humphreys, B. D. Cell profiling of mouse acute kidney injury reveals conserved cellular responses to injury. Proc Natl Acad Sci U S A 117, 15874–15883, doi:10.1073/pnas.2005477117 (2020).

35 Sztalryd, C. & Brasaemle, D. L. The perilipin family of lipid droplet proteins: Gatekeepers of intracellular lipolysis. Biochim Biophys Acta Mol Cell Biol Lipids 1862, 1221–1232, doi:10.1016/j.bbalip.2017.07.009 (2017).

36 Qian, A., Hsieh, T. B., Hossain, M. M., Lin, J. J. & Jin, J. P. A rapid degradation of calponin 2 is required for cytokinesis. Am J Physiol Cell Physiol 321, C355–C368, doi:10.1152/ajpcell.00569.2020 (2021).

37 Kravic, B. et al. Ubiquitin profiling of lysophagy identifies actin stabilizer CNN2 as a target of VCP/p97 and uncovers a link to HSPB1. Mol Cell 82, 2633–2649 e2637, doi:10.1016/j.molcel.2022.06.012 (2022).

38 Faubert, B., Solmonson, A. & DeBerardinis, R. J. Metabolic reprogramming and cancer progression. Science 368, doi:10.1126/science.aaw5473 (2020).

39 Park, J. S. et al. Mechanical regulation of glycolysis via cytoskeleton architecture. Nature 578, 621–626, doi:10.1038/s41586-020-1998-1 (2020).

40 Poglazov, B. F. & Livanova, N. B. Interaction of actin with the enzymes of carbohydrate metabolism. Adv Enzyme Regul 25, 297–305, doi:10.1016/0065-2571(86)90020-8 (1986).

41 Gewin, L. S. Sugar or Fat? Renal Tubular Metabolism Reviewed in Health and Disease. Nutrients 13, doi:10.3390/nu13051580 (2021).

42 Gewin, L., Zent, R. & Pozzi, A. Progression of chronic kidney disease: too much cellular talk causes damage. Kidney Int 91, 552–560, doi:10.1016/j.kint.2016.08.025 (2017).

43 Xu, S. et al. Nuclear farnesoid X receptor attenuates acute kidney injury through fatty acid oxidation. Kidney Int, doi:10.1016/j.kint.2022.01.029 (2022).

44 Idowu, T. O. & Parikh, S. M. A new chapter in lipid signaling and kidney fibrosis. Sci Transl Med 14, eadd2826, doi:10.1126/scitranslmed.add2826 (2022).

45 Paterni, I., Granchi, C., Katzenellenbogen, J. A. & Minutolo, F. Estrogen receptors alpha (ERalpha) and beta (ERbeta): subtype-selective ligands and clinical potential. Steroids 90, 13–29, doi:10.1016/j.steroids.2014.06.012 (2014).

46 Piret, S. E. et al. Loss of proximal tubular transcription factor Kruppel-like factor 15 exacerbates kidney injury through loss of fatty acid oxidation. Kidney Int 100, 1250–1267, doi:10.1016/j.kint.2021.08.031 (2021).

47 Liu, H. & Chen, Y. G. The Interplay Between TGF-beta Signaling and Cell Metabolism. Front Cell Dev Biol 10, 846723, doi:10.3389/fcell.2022.846723 (2022).

48 Hao, J. W. et al. CD36 facilitates fatty acid uptake by dynamic palmitoylation-regulated endocytosis. Nat Commun 11, 4765, doi:10.1038/s41467-020-18565-8 (2020).

