## Supplemental Materials for "Calponin 2 harnesses metabolic reprogramming to determine kidney fibrosis"

### Supplementary Material

### Detailed Methods

#### Mouse models of CKD

Male Balb/c mice weighing approximately 20-22 g were obtained from the Jackson Laboratory (Bar Harbor, Maine). Three mouse CKD models were constructed by employing ischemia-reperfusion injury (IRI), unilateral ureteral obstruction (UUO), and adriamycin (ADR), respectively. For the IRI model, in brief, the left renal pedicle of the mouse was clipped for 35 minutes using microaneurysm clamps. During the ischemic period, the mouse's body temperature was maintained between 36°C and 37.5°C by using a temperature-controlled heating system. At 10 days after IRI, the contralateral kidney was removed, and the mice were then sacrificed at 7 days after nephrectomy. The UUO model was constructed using routine methods as previously described.[S1] The ADR model involved a single intraperitoneal injection of Adriamycin at 15 mg/kg body weight and then sacrificed 21 days after injection.[S2]

Customized short hairpin RNA (ShRNA) specific for the mouse CNN2 gene was ordered from Qianlong Biotech. The target sequence of the murine CNN2 is CTCCAACTTCATCAAGGCCAT. The target sequence of the scrambled ShRNA is CAACAAGATGAAGAGCACCAA. The mouse CNN2-specific ShRNA was subcloned into pLKO.1 vector. The empty vector and ShCNN2 were administered to mice by a hydrodynamic-based gene transfer technique through rapid injection of a large volume of solution through the tail vein as described elsewhere.[S3] Briefly, ShCNN2 (1mg/kg) was diluted in 1.8 ml saline and injected through the tail vein into mouse circulation within 5-10 seconds. Mice from the vehicle group were identically injected with an empty vector. In the IRI model, ShCNN2 was administered respectively at 1 day before unilateral IRI and 1 day before nephrectomy. For the UUO model, ShCNN2 was administered

1 day before and 3 days after the surgery, respectively. In a separate experiment, Etomoxir (HY-50202A, MedChemExpress LLC, Monmouth Junction, NJ) was administrated to mice at a dose of 15 mg/kg body weight 2 days before IRI and nephrectomy, respectively. Serum and kidney samples were collected for further analyses. All proposed animal experiments were approved by the Institutional Animal Care and Use Committee at the University of Connecticut, School of Medicine.

**Human Kidney Biopsy Specimens**

Human kidney biopsy specimen sections and non-tumor kidney tissue sections were obtained from the pathology archive at the University of Pittsburgh Medical Center. Non-tumor kidney tissue samples from the patients who had kidney cell carcinoma and underwent nephrectomy were used as normal controls. All patients included in the presented study have signed the informed consent forms before they underwent kidney biopsy or nephrectomy. All procedures performed in the present study involving human kidney sections were following ethical standards and were approved by the Institutional Review Board at the University of Pittsburgh, School of Medicine. The demographic data are shown in Supplementary Table S1.

**Human serum sample collection and serum metabolomics**

The serum samples used in the current study came from the investigator-initiated clinical trial that was registered in the Chinese Clinical Trial Registry (ChiCTR2000028949). This study was conducted in compliance with the principles of the 1975 Declaration of Helsinki and was approved by the Ethics Committees of the First Affiliated Hospital of Nanjing University of Traditional Chinese Medicine (2019NL-109-02). All participants have signed the consent forms to allow the extra samples to be used for academic purposes. In total, we collected sera from 441 healthy adult volunteers and 388 non-diabetic CKD patients. Inclusion criteria include (1) 20-75 years old; (2) primary kidney disease with a definite diagnosis;

(3) no diabetes history. Serum metabolomics was performed as we previously reported.[S4] The demographic and clinical characteristics of the participants are described in Supplementary Table S2.

**Determination of Serum Creatinine and blood urea nitrogen**

Serum was collected from mice at 18 days after IRI. Blood urea nitrogen (BUN) and serum creatinine (Scr) levels were determined using the QuantiChrom™ Urea (DIUR-100) and Creatinine (DICT-500) assay kits, according to the protocols specified by the manufacturer (BioAssay Systems, Hayward, CA). The levels of BUN and Scr were expressed as milligrams per 100 ml (dL).

**Triglyceride measurement**

The levels of serum and kidney triglyceride were measured by using Triglyceride Quantification Kit (MAK266, Sigma-Aldrich, St. Louis, MO), according to the manufacturer's instructions.

**ATP Measurement**

ATP content in kidney tissue was measured by using the ATP Colorimetric/Fluorometric Assay Kit (K354-100, BioVision, Waltham, MA), according to the manufacturer’s instructions. Data were normalized for total protein content.

**Oil Red O staining**

To detect kidney lipid accumulation, 10 µm thick fresh frozen kidney sections were prepared by a routine procedure. Slides are air-dried for 30 minutes at room temperature and then fixed in ice cold 4% paraformaldehyde. After being placed in 60% isopropanol for 5 minutes, the slides were incubated with freshly prepared Oil Red O (ORO) working solution for 15 minutes. Then, 60% isopropanol was added to re-dissolve the ORO dye. Slides were counter-stained with hematoxylin 3 minutes after ORO staining. Kidney sections were visualized under an Olympus BX43 microscope equipped with a digital camera (Allentown, PA).

**Enzyme-linked immunosorbent assay (ELISA)**

The human Calponin 2 (CNN2, MBS2706690) and rat CNN2 (MBS075726) Elisa kits were purchased from MyBioSource, Inc (San Diego, CA). This assay employs the quantitative sandwich enzyme immunoassay technique. Antibody specific for CNN2 has been pre-coated onto a 96-well strip plate. Prepare 7 wells for standard, 1 well for blank. Add 100 µl of standard, blank, and sample dilutions into the appropriate wells. The plate was then incubated at 37°C for 2 hours. Remove the liquid from each well and directly add 100 µl of Detection Reagent A working solution to each well. The plate was again incubated at 37°C for 1 hour. After washing, add 100 µL of Detection Reagent B working solution to each well and incubated the plate for 30 minutes at 37°C. Following a wash to remove any unbound reagent, 90 µl of substrate solution was added to the wells, and color developed in proportion to the amount of CNN2 bound in the initial step. The color development was stopped, and the intensity of the color was measured immediately using a microplate reader set to 450 nm.

**Quantitative Real-Time Reverse Transcription PCR (qRT-PCR)**

Total RNA isolation and qRT-PCR were carried out by procedures described previously.[S1] Briefly, the first strand cDNA synthesis was carried out using a reverse transcription system kit according to the instructions of the manufacturer (Promega). qRT-PCR was performed on an ABI PRISM 7000 sequence detection system (Applied Biosystems, Foster City, CA). The mRNA levels of various genes were calculated after normalizing with β-actin. Primer sequences used for amplifications are presented in Supplementary Table S3.

### Histology and Immunohistochemical Staining

Paraffin-embedded human kidney biopsy sections (2.5 μm thick) and mouse kidney sections (3 μm thick) were prepared by a routine procedure. The sections were stained with Masson's trichrome reagent (HT15-1KT, Sigma-Aldrich, St. Louis, MO) by standard protocol. Immunohistochemical staining was performed according to the established protocol as described previously.[S1] After incubation with primary antibodies at 4°C overnight, the slides were then stained with HRP-conjugated secondary antibody (Jackson ImmunoResearch Laboratories, West Grove, PA). Non-immune normal IgG was used to replace primary antibodies as a negative control, and no staining was visible. Slides were viewed under an Olympus BX43 microscope equipped with a digital camera (Allentown, PA). The detailed information of the applied primary and secondary antibodies were presented in Supplementary Table S4.

### Immunofluorescence staining

Kidney cryosections were fixed with 3.7% paraformalin for 15 min at room temperature. After blocking with 10% donkey serum for 1 hour, the slides were immunostained with primary antibodies against CD3 and fibronectin. These slides were then stained with Cy2- or Cy3-conjugated secondary antibody (Jackson ImmunoResearch Laboratories, West Grove, PA). Slides were viewed under an Olympus BX43 microscope equipped with a digital camera (Allentown, PA). The detailed information of the applied primary and secondary antibodies were presented in Supplementary Table S4.

### Western Blot Analysis

Kidney tissues were lysed with radioimmune precipitation assay (RIPA) buffer containing 1% NP-40, 0.1% SDS, 100 μg/ml PMSF, 1% protease inhibitor cocktail, and 1% phosphatase I and II inhibitor cocktail (Sigma) in PBS on ice. The supernatants were collected after centrifugation at 13,000×g at 4°C for 15 min. Protein expression was analyzed by western blot as described previously. The detailed information of the applied primary and secondary antibodies were presented in Supplementary Table S4.

**Global Proteomics Sample Preparation**

Kidney tissues were processed for proteomics following a shotgun approach published previously.[S5] In brief, ShNC and ShCNN2 mice kidneys were lysed using SDS buffer (4% SDS, 50mM EDTA, 20mM DTT, 2% Tween 20, 100mM Tris-HCl, pH 8.0) and sonicated (Misonix Sonicator 3000 Ultrasonic Cell Disruptor). The proteins were processed following the suspension trapping (STrap) digestion protocol using in-house packed filters (GF/F, Whatman). The peptides were desalted using the spinnable StageTip protocol, and stored under -80° until further analysis.

**LC-MS/MS analysis**

The LC-MS/MS analysis was performed using an Orbitrap Eclipse MS (Thermo Scientific) coupled with an Ultimate 3000 nanoLC system and a FAIMS Pro Interface (Thermo Scientific). Peptides were first loaded onto a trap column and then separated by an analytical column (PepMap C18, 2.0 um; 15 cm x 75 mm I.D.; Thermo Scientific) at 250 nl/min flow rate using a binary buffer system (buffer A, 0.1% formic acid in water; buffer B, 0.1% formic acid in acetonitrile) with a 165-min gradient (1% to 10% in 8 min; then to 25% buffer B over 117 min; 25% to 32% buffer B in 10 min, then to 95% buffer B over 3 min; back to 1% B in 5 min, and stay equilibration at 1% B for 20 min). Multiple CVs (-40, -60, and -80) were applied for FAIMS separation. For all MS experiments, the survey scans (MS1) were acquired at a resolution of 60,000 in the Orbitrap. The maximum injection time was set to Auto, and AGC target was set to Standard. Monoisotopic peak selection was set to Peptides, and the charge state filter was set to 2-7. For MS/MS acquisition, precursors were isolated with a width of 1.6 m/z, fragmented with HCD using 30% collision energy with a Dynamic maximum injection time, and collected in Orbitrap at 15,000 resolution. The dynamic exclusion was set to 30 s, and can be shared across different FAIMS experiments.

**Protein identification, quantitation, and bioinformatics analysis**

Protein quantitation was performed using the MaxQuant software (version 2.1.3.0) with most of the default parameters, including trypsin as an enzyme with a maximum of two missed cleavage sites; acetylation (protein N-terminal and Lys) and oxidation (Met) as variable modifications; cysteine carbamidomethylation as a fixed modification; peptide length must be at least 7 amino acids; false discovery rate (FDR) were set at 1% for both protein and peptide identification. The MaxQuant output (proteinGroups.txt) was log2-transformed prior to downstream data analyses such as missing value imputation, hierarchical clustering, Principal Component Analysis (PCA), t-tests, correlation, and volcano plots, which were performed in the Perseus environment using default parameters (version 1.6.2.3). Gene Ontology (GO) and Kyoto Encyclopedia of Genes and Genomes (KEGG) pathway analyses were performed using DAVID Bioinformatics Resources 6.8 (https://david.ncifcrf.gov/home.jsp).

**Cell culture and treatment**

Normal rat kidney fibroblasts (NRK-49F) and human kidney proximal tubular cells (HK-2) were obtained from the American Type Culture Collection (ATCC, Manassas, VA). For conditioned media (CM) collections, NRK-49F cells were transfected with siCNN2 (4390815, ThermoFisher Scientific, Waltham, MA) for 24 hours and then cultured with serum-free media for 24 hours. Cultured medium was harvested and centrifuged (3000 rpm for 10 min at 4°C). The conditioned medium (CM) was aliquoted and stored at -80°C for subsequent experiments. Serum-starved HK-2 cells were then transfected with ESR2-Dicer siRNA (hs.Ri.ESR2.13, IDT, Coralville, Iowa) or treated with the CM or CNN2 human recombinant protein (ab177711, abcam, Waltham, MA) or Etomoxir (HY-50202A) or Fenofibrate (HY-17356, MedChemExpress LLC, Monmouth Junction, NJ) under TGFβ1 (240B002; R&D Systems, Minneapolis, MN) stress.

**Chromatin immunoprecipitation (ChIP) Assay**

To analyze the interactions of ESR2 and the binding sites in the promoter of the PPARα gene, a ChIP assay was performed. This assay was carried out according to the protocols specified by the manufacturer (SimpleChIP® Enzymatic Chromatin IP Kit, #9003, Cell Signaling Technology, Danvers, MA). In brief, after various treatments as indicated, HK-2 cells were cross-linked with 1% formaldehyde and then resuspended in SDS lysis buffer containing protease inhibitors. The chromatin solution was fragmented by partial digestion with Micrococcal Nuclease and sonicated, and the supernatant was diluted 10-fold. An aliquot of total diluted lysate was used for total genomic DNA as input DNA control. The anti-ESR2 (GTX70174, GeneTex, Irvine, CA) was added and incubated at 4°C overnight followed by incubation with protein G magnetic beads for 1 hour. The precipitates were washed, and chromatin complexes were eluted. After reversal of the cross-linking at 65°C for 4 hours, the DNA was purified, and ChIP samples were used as a template for qPCR. Primer sets encompass regions of PPARα promoters containing putative binding sites. The sequences of primers used for ChIP assay are given in Supplemental Table S3.

**Automated Analysis of Positive Staining Area in Mouse Kidneys**

Images were randomly selected for each kidney section in the stained slides. The percentage of the positive staining area was analyzed with a custom script in Image Pro plus 6.0, as we previously described.[S1] An average percentage of the positive area for each image was calculated. In this study, the blue channel was chosen because it had the best separation. In brief, by using the "Threshold" tool, the threshold was set to (0, 80) to get the (red) areas where each marker was expressed at a high level. The threshold values were chosen manually until each stained marker was highlighted in red but the same thresholds were used for all images for each marker. In this study, five mice in each group and four randomly selected images *per* mouse were quantified.

**Statistics**

All data were expressed as mean ± SEM if not specified otherwise in the legends. Statistical analysis of the data was performed using GraphPad Prism 9 (GraphPad Software, San Diego, CA). Comparison between two groups was made using a two-tailed Student’s t-test or the Rank Sum Test if data failed a normality test. Statistical significance for multiple groups was assessed by one-way or two-way ANOVA, followed by the Student-Newman-Keuls test. Results are presented in dot plots, with dots denoting individual values. $P < 0.05$ was considered statistically significant.

### References


S1 Fu, H. *et al.* The hepatocyte growth factor/c-met pathway is a key determinant of the fibrotic kidney local microenvironment. *iScience* **24**, 103112, doi:10.1016/j.isci.2021.103112 (2021).

S2 Zhou, D. *et al.* Non-canonical Wnt/calcium signaling is protective against podocyte injury and glomerulosclerosis. *Kidney Int* **102**, 96-107, doi:10.1016/j.kint.2022.02.029 (2022).

S3 Zhou, D. *et al.* Sonic hedgehog is a novel tubule-derived growth factor for interstitial fibroblasts after kidney injury. *J Am Soc Nephrol* **25**, 2187-2200, doi:10.1681/ASN.2013080893 (2014).

S4 Liu, S. *et al.* Serum integrative omics reveals the landscape of human diabetic kidney disease. *Mol Metab* **54**, 101367, doi:10.1016/j.molmet.2021.101367 (2021).

S5 Lin, Y. H. *et al.* Global Proteome and Phosphoproteome Characterization of Sepsis-induced Kidney Injury. *Mol Cell Proteomics* **19**, 2030-2047, doi:10.1074/mcp.RA120.002235 (2020).

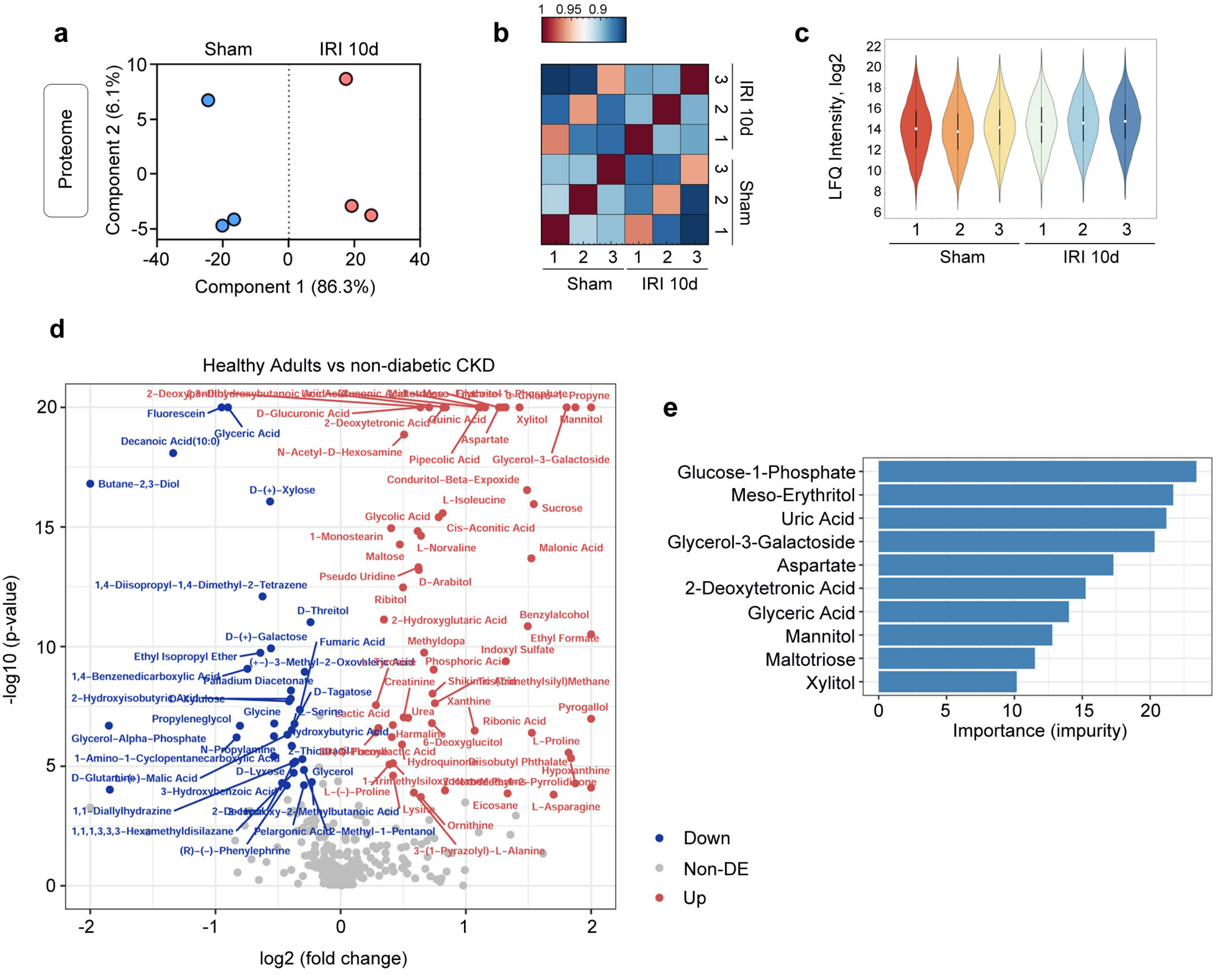


**Supplementary Figure S1. Proteomics and metabolomics profile the landscape of CKD.** (**a**) Principal component analysis of proteomes from control and IRI-induced CKD kidneys. (**b**) Correlation of kidney proteome profiles between control and IRI-induced CKD kidneys. The color scale represents $R^2$ values. (**c**) Violin plot of ANOVA significant proteins (Permutation FDR 0.05) among control and IRI-induced CKD kidneys. LFQ intensities of represented proteins were z-scored and plotted according to the color bar. (**d**) Volcano plot of the differentially expressed serum metabolites for the comparison between healthy adults and non-diabetic CKD patients. (**e**) Top prediction features selected by random forest impurity measurement.

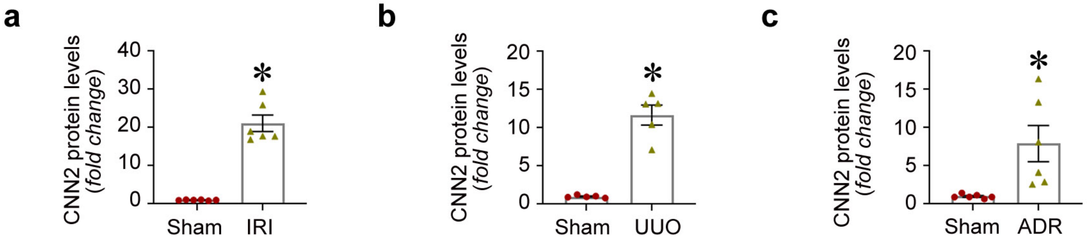


**Supplementary Figure S2: Calponin 2 is upregulated in the fibrotic kidneys after CKD.** (**a-c**) The quantitative data for CNN2 protein expression in the fibrotic kidneys induced by IRI (**a**), UUO (**b**), and ADR nephropathy (**c**). IRI, ischemia reperfusion injury; UUO, unilateral ureteral obstruction; ADR, Adriamycin.

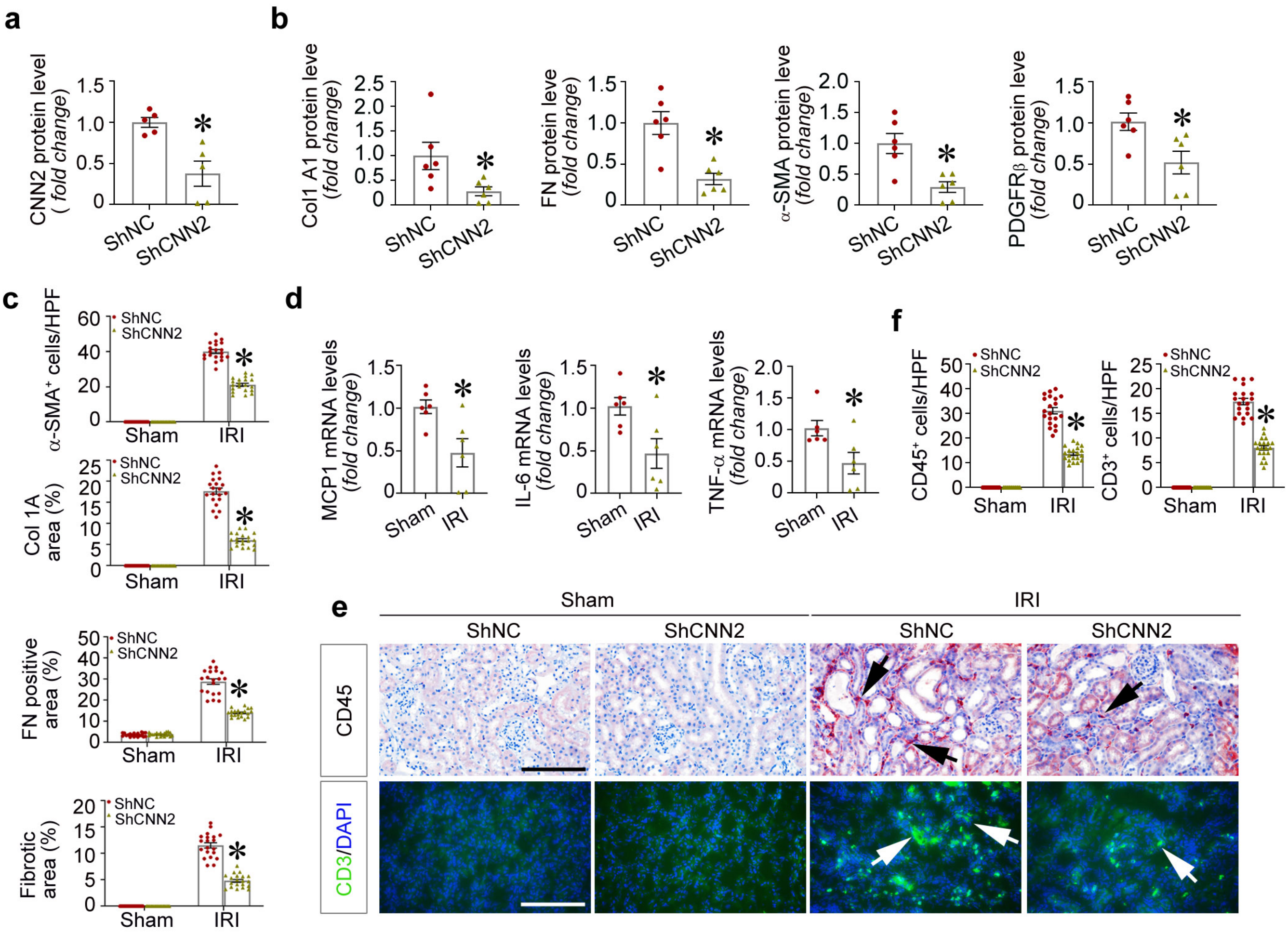


**Supplementary Figure S3. Knockdown of CNN2 alleviates kidney fibrosis induced by IRI.** (**a**) The quantitative data for CNN2 protein expression in the fibrotic kidneys after hydrodynamically delivered control and ShCNN2 plasmid in vivo. Graphs are presented as means ± SEM. * $P < 0.05$ (n=5). (**b**) The quantified data of Col1α1, FN, α-SMA, and PDGFR-β protein expression in ShNC and ShCNN2 mice fibrotic kidneys after IRI-induced CKD. Graphs are presented as means ± SEM. * $P < 0.05$ (n=6). (**c**) The quantitative data for immunohistochemical staining (α-SMA, Col1α1, FN) and Masson Trichrome Staining in ShNC and ShCNN2 mice fibrotic kidneys after IRI-induced CKD. Graphs are presented as means ± SEM. * $P < 0.05$ (n=5, 4 random images were selected in each mouse). (**d**) Quantitative RT-PCR analyses showed the mRNA abundance of MCP-1, IL-6, and TNF-α in ShNC and ShCNN2 mice kidneys after IRI-induced CKD. Graphs are presented as means ± SEM. * $P < 0.05$ (n=6). (**e, f**) Representative micrographs for CD45 and CD3 staining in the fibrotic kidneys collected from ShNC and ShCNN2 mice after IRI-induced CKD (**e**) and quantitative data are presented (**f**). Arrows indicate positive staining. Graphs are presented as means ± SEM. * $P < 0.05$ (n=5). Scale bar, 50 μm. Col1α1, α1 collagen type I; FN, fibronectin; α-SMA, α-smooth muscle actin; PDGFR-β, platelet-derived growth factor receptor β; MCP-1, Monocyte chemoattractant protein-1; IL-6, interleukin 6; TNF-α, tumor necrosis factor-α.

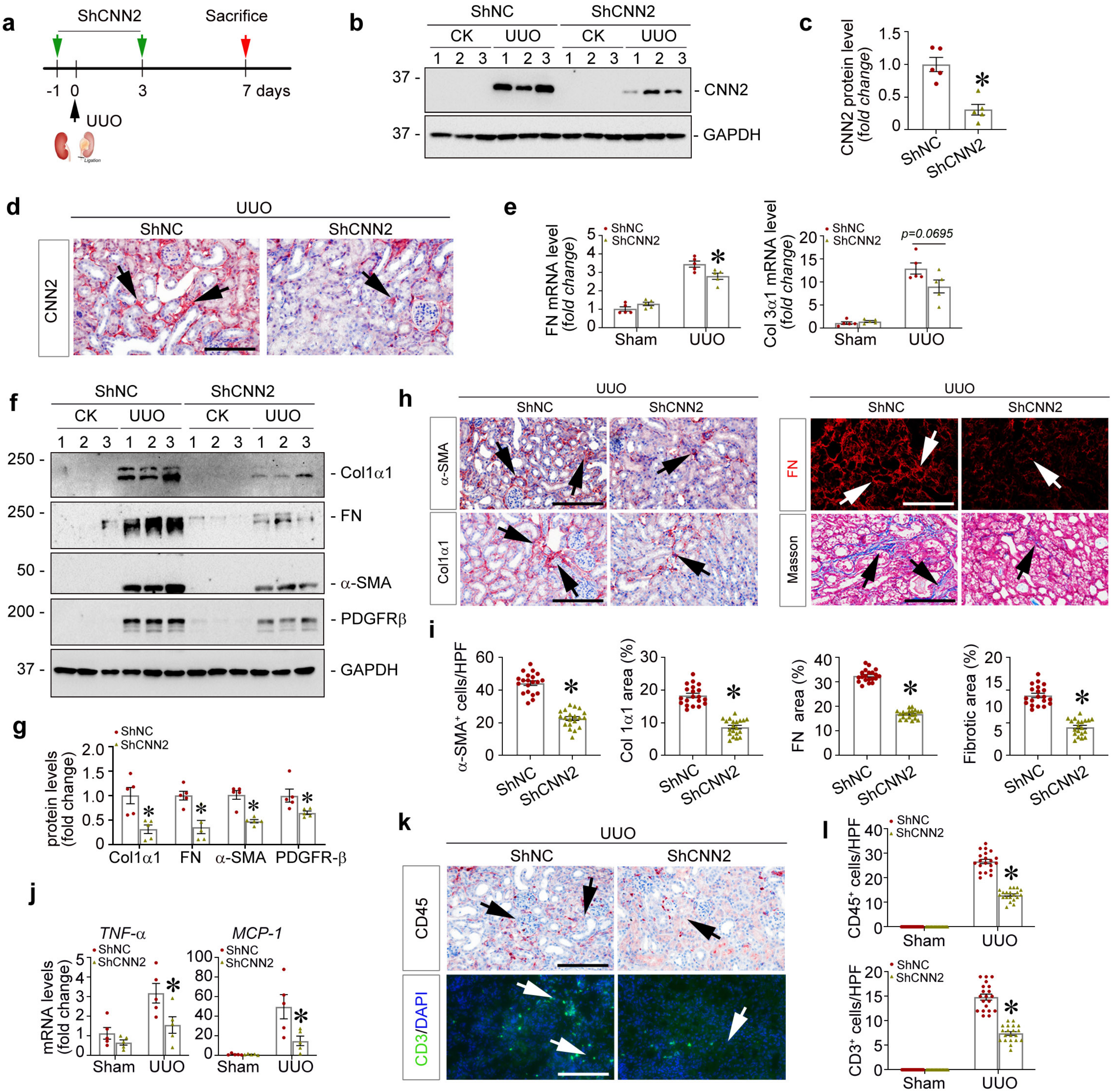

**Supplementary Figure S4: Knockdown of CNN2 ameliorates obstructive CKD in mice.** (**a**) Experiment design. (**b, c**) Western blot assay demonstrated CNN2 protein expression in ShNC and ShCNN2 mice fibrotic kidneys after UUO (**b**) and quantified data are presented (**c**). Numbers indicate individual animals within each group. Graphs are presented as means ± SEM. * *P* <0.05 (n=5). (**d**) Immunohistochemical staining showed CNN2 distributions in ShNC and ShCNN2 mice fibrotic kidneys after UUO. Arrows indicate positive staining. Scale bar, 50 μm. (**e**) Quantitative RT-PCR (qPCR) analyses showed the mRNA abundance of FN and α1 type III collagen (Col3α1) in ShNC and ShCNN2 mice fibrotic kidneys after UUO. Graphs are presented as means ± SEM. * *P* < 0.05 (n=5). (**f, g**) Western blot assays demonstrated α1 type I Collagen (Col1α1), FN, α-SMA, and PDGFRβ protein expression in ShNC and ShCNN2 mice fibrotic kidneys after UUO (**f**) and quantitative data are presented (**g**). Numbers indicate individual animals within each group. Graphs are presented as means ± SEM. * *P* < 0.05 (n=5). (**h, i**) Representative micrographs for α-SMA, Col3α1, FN, and Masson Trichrome staining in ShNC and ShCNN2 mice kidneys after UUO (**h**) and quantitative data are presented (**i**). Arrows indicate positive staining. Graphs are presented as means ± SEM. * *P* <0.05 (n=5). Scale bar, 50 μm. (**j**) qPCR analyses revealed the mRNA abundance of TNF-α and MCP-1 in ShNC and ShCNN2 mice fibrotic kidneys after UUO. Graphs are presented as means ± SEM. * P < 0.05 (n=5). (**k, l**) Representative micrographs for CD45 and CD3 expression in shNC and shCNN2 mice kidneys after UUO (**k**) and quantitative data are presented (**l**). Arrows indicate positive staining. Graphs are presented as means ± SEM. * *P* < 0.05 (n=5). Scale bar, 50 μm.

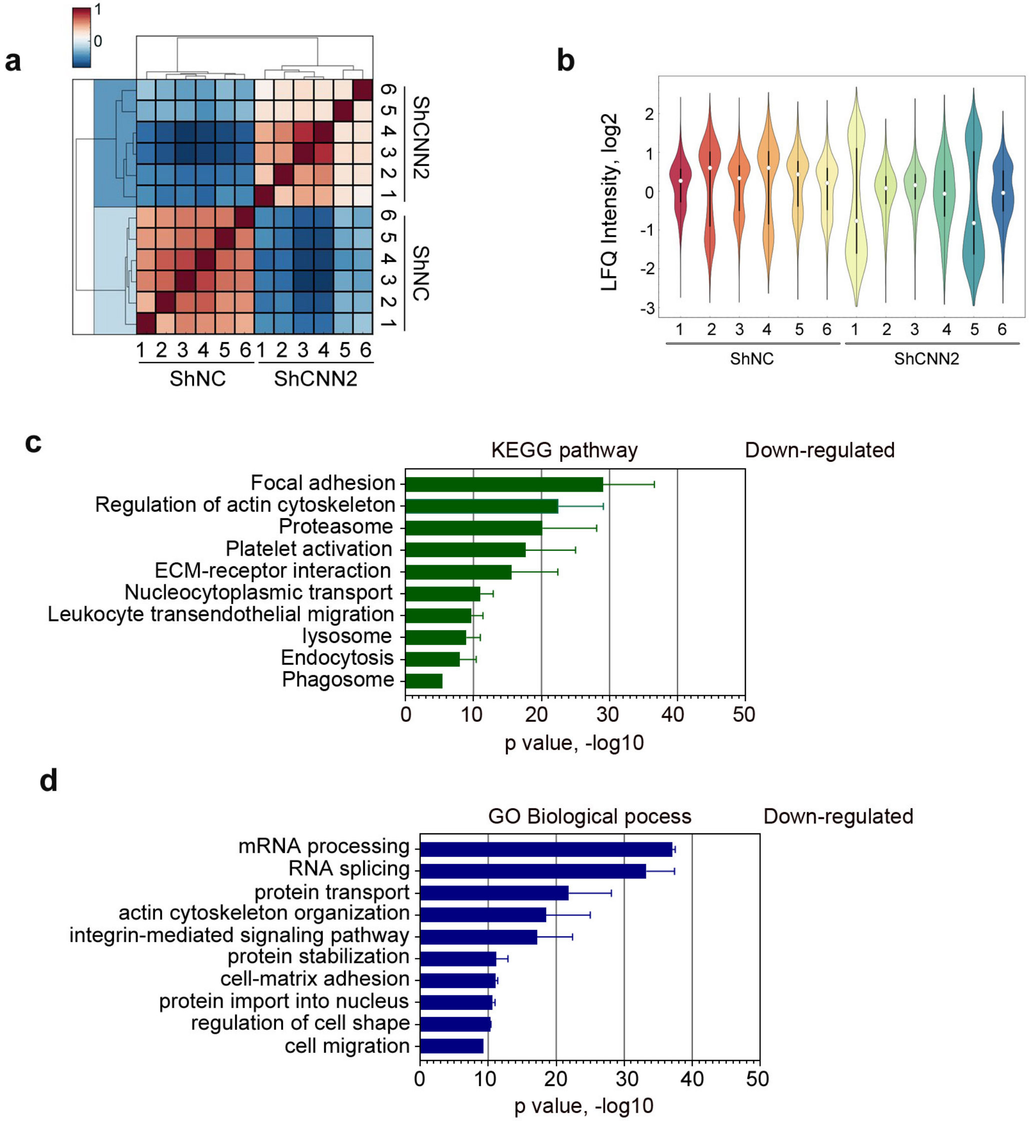


**Supplementary Figure S5: Global proteomics reveals knockdown of CNN2 activates fatty acid oxidation pathway in the fibrotic kidneys after CKD. (a)** Correlation of kidney proteome profiles between ShNC and ShCNN2 mice after IRI-induced CKD. Color scale represents R2 values. (**b**) Violin plot of ANOVA significant proteins (Permutation FDR 0.05) among ShNC and ShCNN2 mice fibrotic kidneys after IRI-induced CKD. LFQ intensities of represented proteins were z-scored and plotted according to the color bar. (**c**) KEGG pathway enrichment analysis showed down-regulated pathway in ShNC and ShCNN2 mice fibrotic kidneys after IRI-induced CKD. (**d**) Gene Ontology (GO) enrichment analysis under biological process terms showed the cluster of down-regulated proteins with their names and significance.

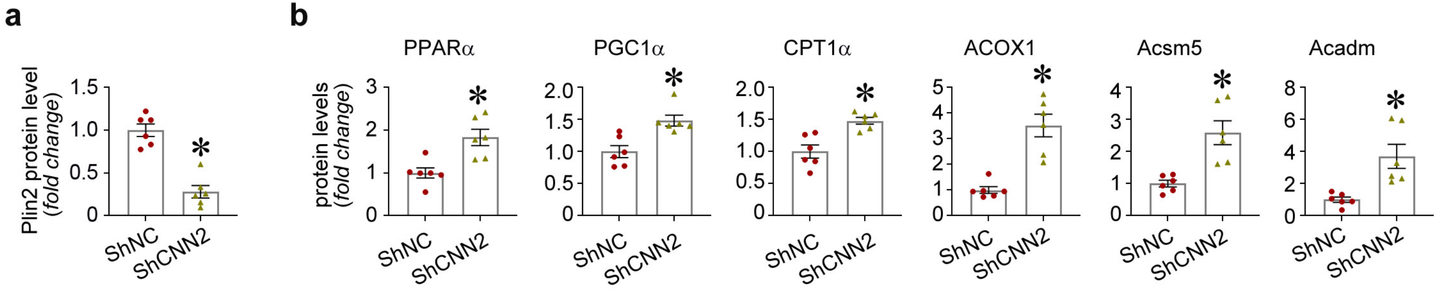


**Supplementary Figure S6: Knockdown of CNN2 decreases lipid accumulation and activates FAO pathway after IRI-induced CKD.**
(**a, b**) The quantitative data for the protein expression of Plin2, PPARα, PGC1α, CPT1α, ACOX1, Acsm5, and Acadm in ShNC and ShCNN2 mice fibrotic kidneys after IRI-induced CKD. Graphs are presented as means ± SEM. * $P < 0.05$ (n=6). Plin2, perilipin 2; PPARα, Peroxisome proliferator-activated receptor α; PGC1α, peroxisome proliferator-activated receptor gamma coactivator 1-α; CPT1α, Carnitine palmitoyltransferase 1α; ACOX1, Acyl-CoA Oxidase 1; Acsm5, Acyl-CoA Synthetase Medium Chain Family Member 5; Acadm, acyl-Coenzyme A dehydrogenase, C-4 to C-12 straight chain.

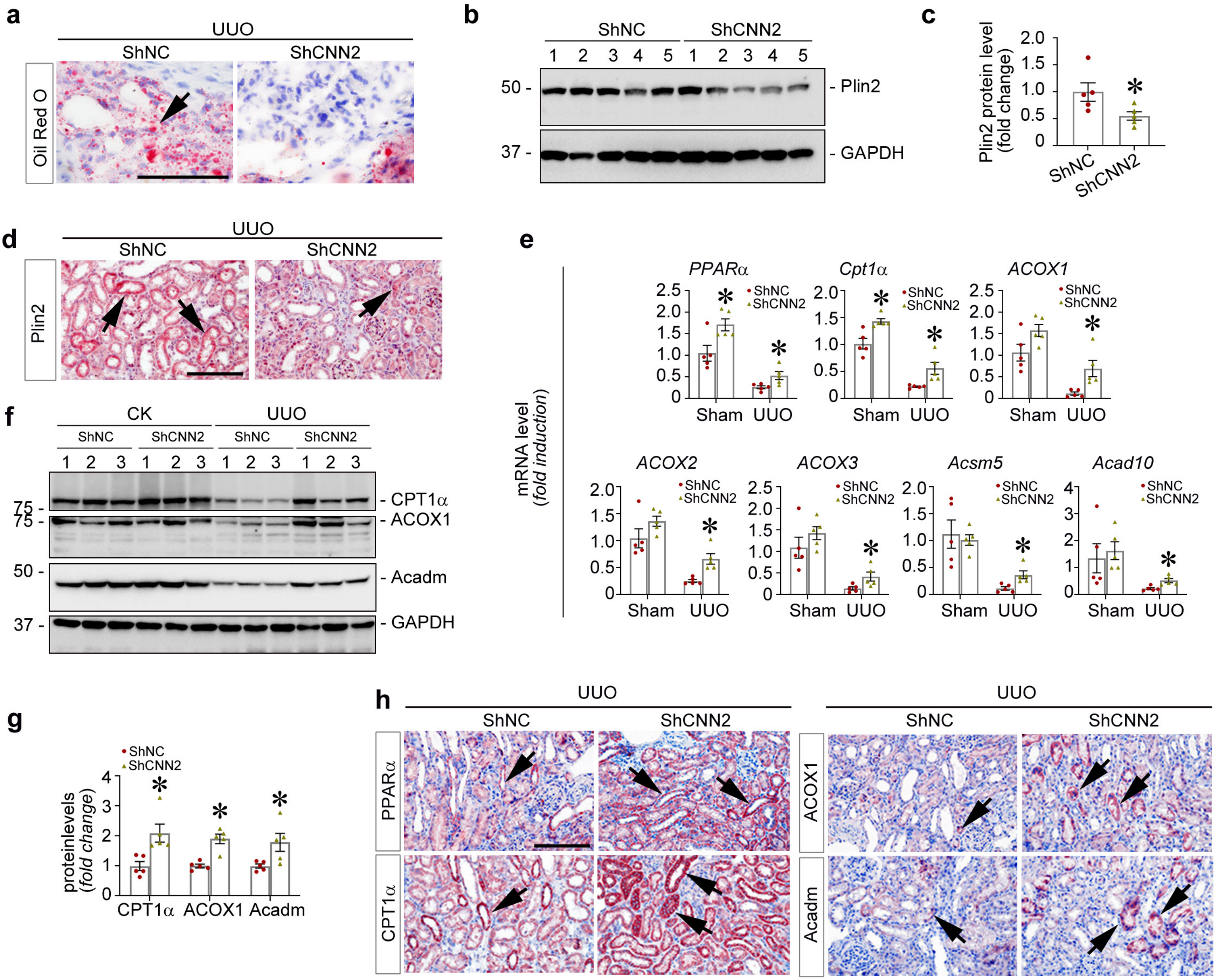


**Supplementary Figure S7: Knockdown of CNN2 decreases lipid accumulation and activates fatty acid oxidation pathway after obstructive CKD.** (**a**) Representative micrographs Oil Red-O staining in the ShNC and ShCNN2 mice fibrotic kidneys after UUO. Arrows indicate the formed lipid droplet. Scale bar, 50 μm. (**b, c**) Western blot analyses demonstrated Plin2 protein expression in ShNC and ShCNN2 mice fibrotic kidneys after UUO (B) and quantitative data are presented (C). Numbers indicate individual animals within each group. Graphs are presented as means ± SEM. * $P < 0.05$ (n=5). (**d**) Representative micrographs Plin2 staining in the kidneys collected from ShNC and ShCNN2 mice after UUO. Arrows indicate positive staining. Scale bar, 50 μm. (**e**) Quantitative RT-PCR analyses showed the mRNA abundance of PPARα, CPT1α, ACOX1, ACOX2, ACOX3, Acsm5, and Acad10 in ShNC and ShCNN2 mice fibrotic kidneys after UUO. Graphs are presented as means ± SEM. * $P < 0.05$ (n=5). (**f, g**) Western blot analyses demonstrated CPT1α, ACOX1, and Acadm protein expression in ShNC and ShCNN2 mice fibrotic kidneys after UUO (**f**) and quantitative data are presented (**g**).Graphs are presented as means ± SEM. * $P < 0.05$ (n=5). (**h**) Representative micrographs PPARα, CPT1α, ACOX1, and Acadm staining in the ShNC and ShCNN2 mice fibrotic kidneys after UUO. Scale bar, 50 μm. Arrows indicate positive staining. Plin2, perilipin 2; PPARα, Peroxisome proliferator-activated receptor α; PGC1α, peroxisome proliferator-activated receptor gamma coactivator 1-α; CPT1α, Carnitine palmitoyltransferase 1α; ACOX1, 2, 3, Acyl-CoA Oxidase 1, 2, 3; Acsm5, Acyl-CoA Synthetase Medium Chain Family Member 5; Acad10, Acyl-CoA Dehydrogenase Family Member 10. Acadm, acyl-Coenzyme A dehydrogenase, C-4 to C-12 straight chain.

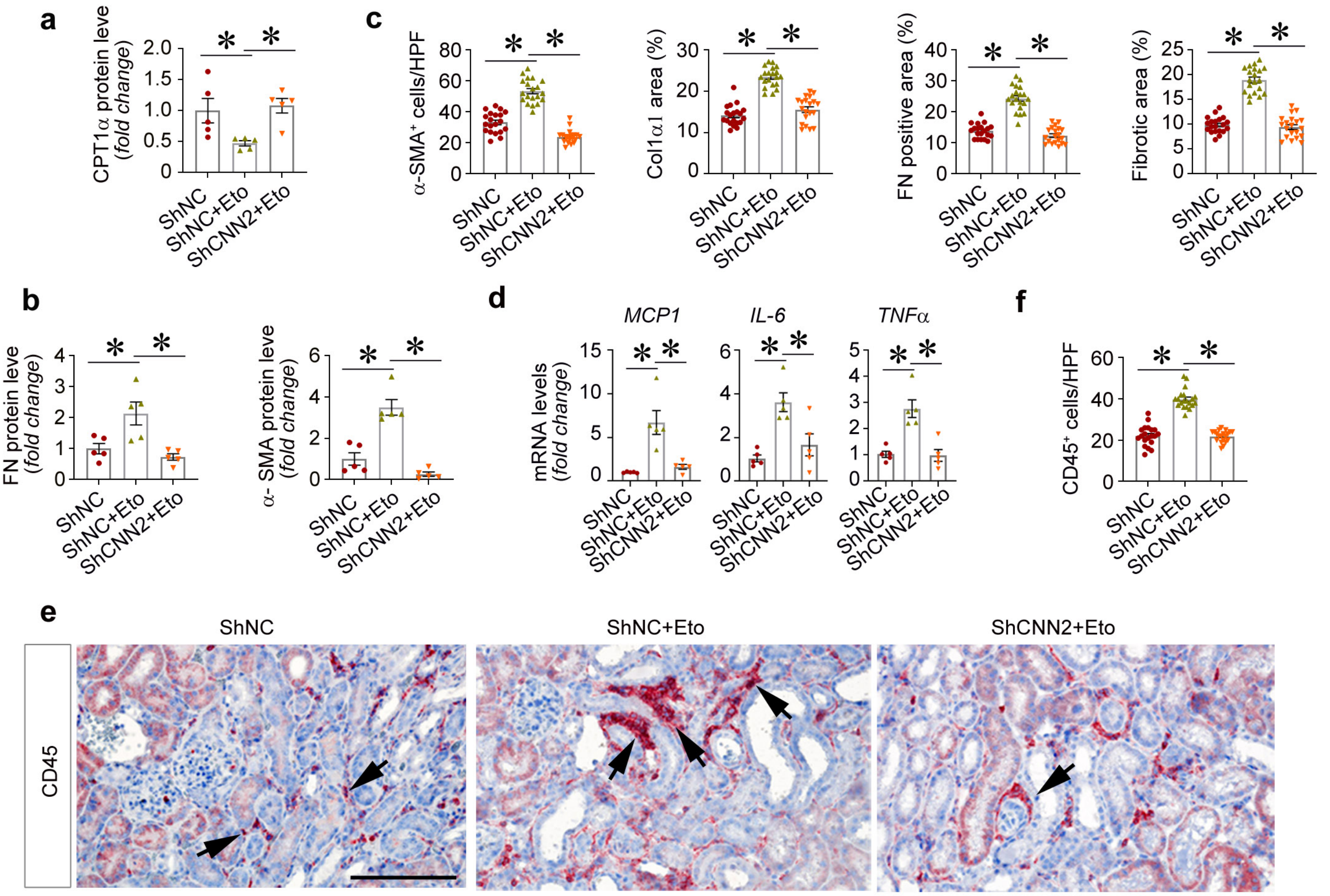


**Supplementary Figure S8. Knockdown of CNN2 enhances CPT1α activity to alleviate kidney fibrosis** (**a, b**) The quantitative data for CPT1α, FN, and α-SMA protein expression in the ShNC mice, ShNC mice received Etomoxir, and ShCNN2 mice received Etomoxir fibrotic kidneys after IRI-induced CKD. Graphs are presented as means ± SEM. * P < 0.05 (n=5). (**c**) The quantitative data for α-SMA, Col1α1, FN, and Masson Trichrome Staining in the fibrotic kidneys collected from ShNC mice, ShNC mice received Etomoxir, and ShCNN2 mice received Etomoxir after IRI-induced CKD. Graphs are presented as means ± SEM. * P < 0.05 (n=5). (**d**) Quantitative RT-PCR analyses showed the mRNA abundance of MCP-1, IL-6, and TNF-α in ShNC mice, ShNC mice received Etomoxir, and ShCNN2 mice received Etomoxir fibrotic kidneys after IRI-induced CKD. Graphs are presented as means ± SEM. * P < 0.05 (n=5). (**e, f**) Representative micrographs for immunostaining of CD45 showed inflammatory cells infiltration in the fibrotic kidneys collected from ShNC mice, ShNC mice received Etomoxir, and ShCNN2 mice received Etomoxir after IRI-induced CKD (**e**) and the quantitative data are presented (**f**). Arrows indicate positive staining. Scale bar, 50 μm.

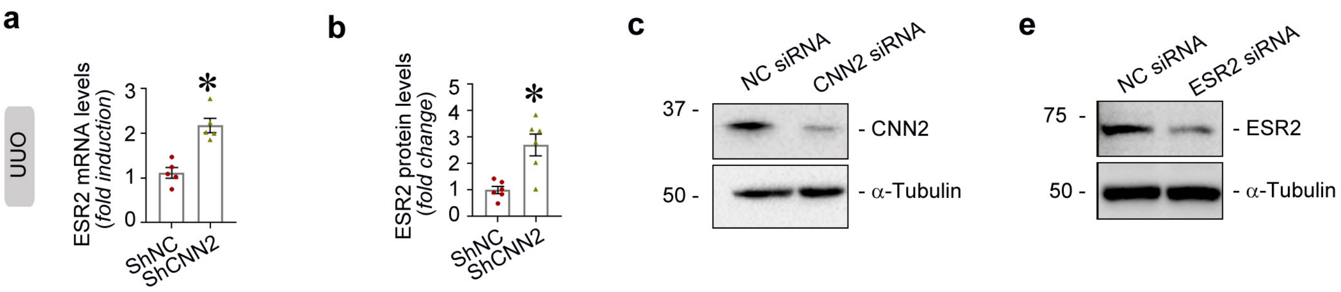


d

| Matrix ID | Name | Score | Relative score | Sequence ID | Start | End | Strand | Predicted sequence |
|---|---|---|---|---|---|---|---|---|
| MA0258.2 | MA0258.2. ESR2 | 12.748039 | 0.898178519 | NC_000022.11:46148526-46150525 | 1274 | 1288 | + | AGGGCACCCGGTCCC |
| MA0258.2 | MA0258.2. ESR2 | 10.833046 | 0.872880315 | NC_000022.11:46148526-46150525 | 570 | 584 | + | GGGTCAGCTTCCCTC |
| MA0258.2 | MA0258.2. ESR2 | 8.3418865 | 0.839970612 | NC_000022.11:46148526-46150525 | 661 | 675 | + | AGGTCACCCTATAAA |
| MA0258.2 | MA0258.2. ESR2 | 8.192669 | 0.837999358 | NC_000022.11:46148526-46150525 | 54 | 68 | + | AGGTCTTCGTGGCCA |
| MA0258.2 | MA0258.2. ESR2 | 8.088699 | 0.836625858 | NC_000022.11:46148526-46150525 | 3 | 17 | + | GGGTGAGGGTGTGCC |
| MA0258.2 | MA0258.2. ESR2 | 7.226689 | 0.825238185 | NC_000022.11:46148526-46150525 | 54 | 68 | - | TGGCCACGAAGACCT |
| MA0258.2 | MA0258.2. ESR2 | 6.6058273 | 0.817036233 | NC_000022.11:46148526-46150525 | 5 | 19 | - | AGGGCACACCCTCAC |
| MA0258.2 | MA0258.2. ESR2 | 6.602616 | 0.816993807 | NC_000022.11:46148526-46150525 | 1222 | 1236 | - | GAGGGACCTTGTCCC |
| MA0258.2 | MA0258.2. ESR2 | 5.934113 | 0.808162486 | NC_000022.11:46148526-46150525 | 726 | 740 | + | AGGTGAGGTTGCCGT |
| MA0258.2 | MA0258.2. ESR2 | 5.802187 | 0.806419664 | NC_000022.11:46148526-46150525 | 345 | 359 | - | TGGTCACGTACACTC |
| MA0258.2 | MA0258.2. ESR2 | 5.6550326 | 0.804475668 | NC_000022.11:46148526-46150525 | 1184 | 1198 | + | AAGTCAGCCTTCTCC |
| MA0258.2 | MA0258.2. ESR2 | 5.3605804 | 0.800585779 | NC_000022.11:46148526-46150525 | 218 | 232 | - | CTGTCAGCCAGTCTC |

**Supplementary Figure S9: Knockdown of CNN2 promotes ESR2 binds to PPARα.** (**a**) Quantitative RT-PCR (qPCR) analysis showed the abundance of ESR2 mRNA levels in ShNC and ShCNN2 mice fibrotic kidneys after UUO-induced CKD. Graphs are presented as means ± SEM. * $P$ <0.05 (n=5). (**b**) The quantitative data for ESR2 protein expression after IRI-induced CKD. Graphs are presented as means ± SEM. * $P$ <0.05 (n=6). (**c**) Western blot assay demonstrated the knockdown of CNN2 in normal rat kidney fibroblasts (NRK-49F). (**d**)Through the JASPAR database (https://jaspar.genereg.net/), 12 putative binding sites were identified for ESR2 on the PPARα gene promoter region. (**e**) Western blot assay demonstrated the knockdown of ESR2 in human kidney proximal tubular cells (HK-2).

**Supplementary Table S1. The demographic and clinical data of the patients (Kidney biopsy).**

| No. | Gender | Age | Diagnosis | UTP (g/24h) | Albumin (g/L) | Scr (μmol/L) | BUN (mmol/L) |
|---|---|---|---|---|---|---|---|
| 1 | Female | 60 | FSGS | 4.303 | 15.5 | 65.4 | 3.99 |
| 2 | Female | 51 | FSGS | 5.873 | 23.8 | 66.6 | 8.16 |
| 3 | Male | 49 | FSGS | 7.835 | 17.7 | 124.9 | 12.94 |
| 4 | Male | 55 | FSGS | 7.916 | 23.1 | 153.9 | 11.32 |
| 5 | Male | 47 | FSGS | 3.227 | 27.1 | 102.4 | 9.16 |
| 6 | Female | 34 | IgAN | 4.429 | 30.2 | 86.5 | 5.28 |
| 7 | Male | 39 | IgAN | 6.422 | 18.9 | 108 | 5.13 |
| 8 | Male | 36 | IgAN | 17.97 | 39 | 171 | 8.96 |
| 9 | Male | 25 | IgAN | 11.54 | 17.1 | 96 | 7.45 |
| 10 | Female | 51 | IgAN | 5.421 | 30.4 | 132.9 | 8.07 |
| 11 | Female | 44 | MN | 11.561 | 20.6 | 64.6 | 6.77 |
| 12 | Female | 46 | MN | 3.758 | 25.1 | 64.2 | 5.88 |
| 13 | Female | 29 | MN | 8.253 | 16.9 | 51.5 | 5.42 |
| 14 | Male | 17 | MN | 7.338 | 24.1 | 75.3 | 5.35 |
| 15 | Male | 46 | MN | 12.301 | 22.9 | 87.4 | 6.92 |

FSGS, focal segmental glomerulosclerosis; IgAN, IgA nephropathy; MN, membranous nephropathy; BUN, blood urea nitrogen; Scr, serum creatinine; UTP, urinary total protein.

**Supplementary Table S2. The demographic and clinical data of the participants (metabolomics)**

| **Health Adults** | | | | | |
|---|---|---|---|---|---|
| | Minimum | Maximum | Mean | Standard deviation | Count |
| Age | 20 | 75 | 33.24 | 9.08 | 441 |
| Gender | 259 (Female) | 182 (Male) | | | 441 |
| **Non-Diabetic CKD Patients** | | | | | |
| Age | 19 | 75 | 52.17 | 14.3 | 338 |
| Gender | 157 (Female) | 181 (Male) | | | 338 |
| HbA1c | 4.4 | 10 | 6.02 | 0.94 | 257 |
| eGFR | 2.36 | 143.56 | 42.35 | 36.31 | 335 |
| Albumin | 4.65 | 48 | 31.9 | 7.24 | 334 |
| BUN | 2.97 | 72.2 | 16.13 | 10.97 | 332 |
| Scr | 38.3 | 1970.2 | 324.26 | 306.68 | 335 |
| Uric Acid | 105.7 | 919.9 | 430.64 | 132.39 | 333 |
| TC | 1.8 | 15.56 | 4.96 | 1.88 | 327 |
| TG | 0.25 | 11.01 | 2.02 | 1.35 | 327 |
| HDL-C | 0.44 | 2.85 | 1.29 | 0.39 | 326 |
| LDL-C | 0.84 | 8.36 | 2.93 | 1.28 | 326 |
| CO2CP | 8.9 | 34 | 21.62 | 3.84 | 327 |
| UTP | 2.12 | 19180.6 | 3009.78 | 3482.41 | 297 |

HbA1c, hemoglobin A1C; eGFR, estimated Glomerular Filtration Rate; BUN, blood urea nitrogen; Scr, serum creatinine; TC, total cholesterol; TG, triglycerides; HDL-C, high-density lipoprotein cholesterol; LDL-C, low-density lipoprotein cholesterol; CO2CP, carbon dioxide combining power; UTP, urinary total protein.

**Supplementary Table S3. Nucleotide sequences of the primers used for qRT-PCR**

| gene | Primer Sequence 5' to 3' | |
|---|---|---|
| | Forward | Reverse |
| CNN2 (M) | AGGAAGCAGAACTCCGAAGC | CCAGTTCTGCATAGAGCGGT |
| FN (M) | CGAGGTGACAGAGACCACAA | CTGGAGTCAAGCCAGACACA |
| Col1α1(M) | ATCTCCTGGTGCTGATGGAC | ACCTTGTTTGCCAGGTTCAC |
| Col3α1 (M) | AGGCAACAGTGGTTCTCCTG | GACCTCGTGCTCCAGTTAGC |
| PPARα (M) | ACCACTACGGAGTTCACGCATG | GAATCTTGCAGCTCCGATCACAC |
| CPT1α (M) | GGCATAAACGCAGAGCATTCCTG | CAGTGTCCATCCTCTGAGTAGC |
| ACOX1 (M) | GCCATTCGATACAGTGCTGTGAG | CCGAGAAAGTGGAAGGCATAGG |
| ACOX2 (M) | CAATGGCTTCCTGCGACTGAAC | AAGCCTCTGGTAGGTGCCATCT |
| ACOX3 (M) | CCTATGCCTTGGACCACTTCTC | ATGCCAGAGCATGGATCTCACG |
| Acsm1 (M) | CAGTGGAAGCAAAGGACAGGTC | TCCCTATGGAGCCTCGCTTGAT |
| Acsm2 (M) | CTGGAAAGCCAAGACAGGACTG | TGAGCAAAGGCTGTTCCCAGGT |
| Acsm5 (M) | TCTTCTCTGCCTGGTCCAATGG | AAGAGGGTTGGGACACAGCACA |
| Acadm (M) | AGGATGACGGAGCAGCCAATGA | GCCGTTGATAACATACTCGTCAC |
| Acadl (M) | GGCGATTTCTGCCTGTGAGTTC | GCTGTCCACAAAAGCTCTGGTG |
| Acad10 (M) | GGAACTTTCCACCTTAGGAGACC | CAGCTTTGTCACATCCTGGTCTC |
| MCP1 (M) | TTAAAAACCTGGATCGGAACCAA | GCATTAGCTTCAGATTTACGGGT |
| ESR2 (M) | GGTCCTGTGAAGGATGTAAGGC | TAACACTTGCGAAGTCGGCAGG |
| IL-6 (M) | CTTGGGACTGATGCTGGTG | TCCACGATTTCCCAGAGAAC |
| TNFα (M) | CCCTCACACTCAGATCATCTTCT | CCCTCACACTCAGATCATCTTCT |
| β-actin (M) | CAGCTGAGAGGGAAATCGTG | CGTTGCCAATAGTGATGACC |
| PGC1α (H) | CCAAAGGATGCGCTCTCGTTCA | CGGTGTCTGTAGTGGCTTGACT |
| CPT1α (H) | GATCCTGGACAATACCTCGGAG | CTCCACAGCATCAAGAGACTGC |
| PPARα (H) | TCGGCGAGGATAGTTCTGGAAG | GACCACAGGATAAGTCACCGAG |
| ACOX1 (H) | GGCGCATACATGAAGGAGACCT | AGGTGAAAGCCTTCAGTCCAGC |
| Acsm1 (H) | CTGCTCTACGAGAACTATGGGC | CTGTGTTAGGTGGCAGGATGCT |
| Acsm5 (H) | CCATCTTTCGGCTGCTTGTGCA | CAGTCTGGTGTTTCCACTTCTCC |
| ESR2 (H) | ATGGAGTCTGGTCGTGTGAAGG | TAACACTTCCGAAGTCGGCAGG |
| GAPDH (H) | GTCTCCTCTGACTTCAACAGCG | ACCACCCTGTTGCTGTAGCCAA |
| PPARα (ChIP) | GGTAATGTCTTTGAGCCCGGA | CATATCTCGTGGGGTGTGCAG |
| PPARα (ChIP) | CACCCTACGCACTTCTGAGC | CCCTCCTGCCTCCTCGAT |

**Supplementary Table S4. The information of the applied primary and secondary antibodies**

| Name | Vendor | Category Number | Application |
|---|---|---|---|
| α-SMA | Abcam, Cambridge, MA | ab5694 | WB, IHC |
| CD3 | Abcam, Cambridge, MA | ab135372 | IF |
| CD45 | Cell signaling Technology, Danvers, MA | #70257 | IHC |
| Col1α1 | Cell signaling Technology, Danvers, MA | #72026 | WB, IHC |
| PDGFR-β | Cell signaling Technology, Danvers, MA | #3169 | WB |
| Plin2 | LsBio, Seattle, WA | LS-B15357 | WB, IHC |
| Acadm | Proteintech Group, Rosemont, IL | 55210-1-AP | WB, IHC |
| ACOX1 | Proteintech Group, Rosemont, IL | 10957-1-AP | WB, IHC |
| Acsm5 | Proteintech Group, Rosemont, IL | 16591-1-AP | WB |
| CNN2 | Proteintech Group, Rosemont, IL | 21073-1-AP | WB, IHC |
| CPT1α | Proteintech Group, Rosemont, IL | 15184-1-AP | WB, IHC |
| ESR2 | Proteintech Group, Rosemont, IL | 14007-1-AP | WB |
| ESR2 | GeneTex, Irvine, CA | GTX70174 | ChIP |
| PGC1α | Proteintech Group, Rosemont, IL | 66369-1-Ig | WB |
| PPARα | Proteintech Group, Rosemont, IL | 15540-1-AP | WB, IHC |
| GAPDH | Santa Cruz Biotechnology, Dallas, Texas | sc-32233 | WB |
| Fibronectin | Sigma, St. Louis, MO | F3648 | WB, IF |
| α-Tubulin | Sigma, St. Louis, MO | T9026 | WB |
| Anti-Rabbit IgG | abcam | ab6721 | WB |
| Anti-Mouse IgG | abcam | ab6789 | WB |
| Cy3 Anti-Rabbit | Jackson ImmunoResearch | 711-165-152 | IF |
| Alexa Fluor® 488 Anti-Rabbit | Jackson ImmunoResearch | 711-545-152 | IF |
| Biotin-Anti-Rabbit | Jackson ImmunoResearch | 711-065-152 | IHC |

WB, western blot; IHC, Immunohistochemical staining; IF, Immunofluorescence staining; ChIP, Chromatin immunoprecipitation.

### Supplementary Table S5. Differentially expressed metabolites in health adults and CKD patients

| Metabolite | Denominator | Numeritor | Log2FC | P Value | adj_pValue |
|---|---|---|---|---|---|
| (+-)-3-Methyl-2-Oxovaleric Acid | 15.35455933 | 15.06773948 | -0.286819842 | 1.1367E-09 | 3.45556E-07 |
| (R)-(-)-Phenylephrine | 25.56044722 | 25.12995945 | -0.430487762 | 6.35333E-05 | 0.016264519 |
| 1,1,1,3,3,3-Hexamethyldisilazane | 14.67523315 | 14.30067982 | -0.374553332 | 1.89E-05 | 0.00498959 |
| 1,1-Diallylhydrazine | 14.17296414 | 13.8109348 | -0.362029341 | 6.4405E-06 | 0.001738935 |
| 1,4-Benzenedicarboxylic Acid | 18.11693376 | 17.37247431 | -0.74445945 | 8.49504E-10 | 2.59948E-07 |
| 1,4-Diisopropyl-1,4-Dimethyl-2-Tetrazene | 14.6685303 | 14.04474426 | -0.62378604 | 7.99769E-13 | 2.51927E-10 |
| 1-Amino-1-Cyclopentanecarboxylic Acid | 15.45463885 | 14.62344199 | -0.831196862 | 6.26038E-07 | 0.000174038 |
| 1-Methyl-2-Pyrrolidinone | 8.493421776 | 11.47265663 | 2.979234857 | 7.86307E-05 | 0.020050816 |
| 1-Monostearin | 13.07425638 | 13.4772772 | 0.403020814 | 1.12353E-15 | 3.62899E-13 |
| 1-Trimethylsiloxyoctane | 14.57840609 | 14.99594136 | 0.417535271 | 7.50474E-06 | 0.002018774 |
| 2,3-Dihydroxybutanoic Acid | 18.08143234 | 18.92082605 | 0.83939371 | 3.78696E-22 | 1.26106E-19 |
| 2-Decanol | 15.37330212 | 14.90567047 | -0.46763165 | 5.02318E-05 | 0.013060256 |
| 2-Deoxypentitol | 14.53288694 | 15.23963432 | 0.706747384 | 2.2716E-27 | 7.65528E-25 |
| 2-Deoxytetronic Acid | 17.6705035 | 18.49293077 | 0.822427275 | 7.73845E-35 | 2.63107E-32 |
| 2-Hydroxy-2-Methylbutanoic Acid | 20.17854689 | 19.94815384 | -0.230393052 | 4.62478E-05 | 0.012070682 |
| 2-Hydroxyglutaric Acid | 15.75308862 | 16.09869499 | 0.345606372 | 7.4575E-12 | 2.34165E-09 |
| 2-Hydroxyisobutyric Acid | 25.98637832 | 25.5741875 | -0.412190813 | 1.93371E-08 | 5.80113E-06 |
| 2-Methyl-1-Pentanol | 13.98504966 | 13.69187715 | -0.293172513 | 1.43848E-05 | 0.003811984 |
| 2-Thiouracil | 11.55911147 | 11.17029749 | -0.388813978 | 1.45328E-06 | 0.000399653 |
| 3-(1-Pyrazolyl)-L-Alanine | 16.10568043 | 16.69067498 | 0.584994547 | 0.000126987 | 0.03200077 |
| 3-Chloro-1-Propyne | 8.979234584 | 14.96625648 | 5.987021895 | 1.00656E-23 | 3.37199E-21 |
| 3-Hydroxybenzoic Acid | 17.42181102 | 17.04211379 | -0.379697224 | 7.64962E-06 | 0.002050097 |
| 6-Deoxyglucitol | 14.51141277 | 15.24015787 | 0.728745105 | 1.58338E-07 | 4.62347E-05 |
| Aspartate | 13.84959977 | 15.11397364 | 1.264373873 | 1.609E-42 | 5.53497E-40 |
| Benzylalcohol | 14.62201921 | 16.1162788 | 1.494259587 | 1.40526E-11 | 4.38442E-09 |
| Butane-2,3-Diol | 14.64391657 | 12.08192981 | -2.561986752 | 1.55677E-17 | 5.12177E-15 |
| Cis-Aconitic Acid | 13.292061 | 14.07461195 | 0.782550951 | 3.89764E-16 | 1.26283E-13 |
| Conduritol-Beta-Expoxide | 17.10282909 | 18.59155002 | 1.488720932 | 2.87019E-17 | 9.41423E-15 |
| Creatinine | 18.62339998 | 19.12691456 | 0.50351458 | 9.038E-08 | 2.66621E-05 |
| D-(+)-Fucose | 17.28184595 | 17.56159849 | 0.279752539 | 3.93661E-07 | 0.000111406 |
| D-(+)-Galactose | 26.67759078 | 26.12089815 | -0.556692635 | 1.18363E-10 | 3.66925E-08 |
| D-(+)-Xylose | 20.01725355 | 19.45357608 | -0.563677473 | 8.60735E-17 | 2.8146E-14 |
| D-3-Phenyllactic Acid | 14.83614433 | 15.24404467 | 0.407900337 | 6.04564E-07 | 0.000168673 |
| D-Arabitol | 19.22452239 | 19.84836731 | 0.623844922 | 6.25159E-14 | 1.98175E-11 |
| Decanoic Acid(10:0) | 12.74738838 | 11.40968539 | -1.337702995 | 8.17064E-19 | 2.69631E-16 |
| D-Glucuronic Acid | 17.55935488 | 18.19533651 | 0.635981634 | 2.27301E-22 | 7.59187E-20 |
| D-Glutamine | 17.90409784 | 16.06051884 | -1.843578999 | 9.55493E-05 | 0.024269513 |
| Diisobutyl Phthalate | 13.8228146 | 15.69716978 | 1.874355181 | 5.241E-05 | 0.013574197 |
| D-Lyxose | 18.31279081 | 17.78031627 | -0.532474535 | 3.86768E-06 | 0.001055876 |
| D-Tagatose | 21.04569982 | 20.67703632 | -0.3686635 | 1.7034E-07 | 4.93986E-05 |
| D-Threitol | 17.34157416 | 17.10082497 | -0.24074919 | 9.58368E-12 | 2.99969E-09 |
| D-Xylulose | 19.47172985 | 19.07288094 | -0.398848906 | 1.52015E-08 | 4.57567E-06 |
| Eicosane | 15.10641651 | 16.4396508 | 1.333234292 | 0.000136609 | 0.034288811 |
| Ethyl Formate | 11.67604844 | 14.27731876 | 2.601270319 | 3.07905E-11 | 9.57583E-09 |
| Ethyl Isopropyl Ether | 15.78577969 | 15.14410491 | -0.641674773 | 1.82132E-10 | 5.60966E-08 |
| Fluorescein | 18.14245231 | 17.19176465 | -0.950687657 | 7.53517E-22 | 2.50168E-19 |
| Fumaric Acid | 14.50986158 | 14.18411709 | -0.325744491 | 4.36257E-08 | 1.29568E-05 |
| Gluconic Acid | 17.94615555 | 19.10011498 | 1.153959428 | 8.97719E-24 | 3.01634E-21 |
| Glucose-1-Phosphate | 14.90196158 | 16.21924109 | 1.317279509 | 5.96601E-46 | 2.07021E-43 |
| Glyceric Acid | 20.09874583 | 19.19869817 | -0.90004766 | 3.19739E-38 | 1.09351E-35 |
| Glycerol | 23.99760732 | 23.69220611 | -0.305401206 | 5.07202E-06 | 0.001374518 |
| Glycerol-3-Galactoside | 14.53500278 | 16.33999641 | 1.804993637 | 1.6417E-46 | 5.71313E-44 |
| Glycerol-Alpha-Phosphate | 15.48398163 | 13.63281122 | -1.851170404 | 2.01798E-07 | 5.81179E-05 |
| Glycine | 21.86184941 | 21.33168856 | -0.530160856 | 1.6435E-07 | 4.78258E-05 |
| Glycolic Acid | 18.56701355 | 19.18263305 | 0.615619495 | 1.49287E-15 | 4.80703E-13 |
| Harmaline | 16.28103581 | 16.69797422 | 0.416938407 | 1.90666E-07 | 5.51025E-05 |
| Hydroquinone | 13.98060406 | 14.47068087 | 0.490076815 | 1.24262E-06 | 0.000344207 |
| Hydroxybutyric Acid | 22.11733422 | 21.72664584 | -0.390688384 | 1.39824E-06 | 0.000385914 |
| Hypoxanthine | 9.111494739 | 10.95125642 | 1.839761678 | 4.65104E-06 | 0.001265082 |
| Indoxyl Sulfate | 14.8786543 | 16.19735839 | 1.318704094 | 4.10236E-10 | 1.25942E-07 |
| L-(-)-Malic Acid | 16.21070202 | 15.78604808 | -0.424653942 | 4.8145E-07 | 0.000135287 |
| L-(-)-Proline | 19.43971281 | 19.82907287 | 0.389360059 | 8.40695E-06 | 0.002244656 |
| Lactic Acid | 27.96670676 | 28.26839608 | 0.301689326 | 2.53353E-07 | 7.24589E-05 |
| L-Asparagine | 15.92461406 | 17.62552439 | 1.700910336 | 0.000153818 | 0.038454606 |
| L-Isoleucine | 15.89015605 | 16.70477179 | 0.814615744 | 2.64976E-16 | 8.61173E-14 |
| L-Norvaline | 17.22808557 | 17.86976318 | 0.641677614 | 2.33806E-15 | 7.50518E-13 |

| | | | | | |
|---|---|---|---|---|---|
| L-Proline | 12.63444255 | 14.45503528 | 1.820592723 | 2.71461E-06 | 0.000743803 |
| L-Serine | 19.08318443 | 18.69327702 | -0.38990741 | 3.05752E-07 | 8.71394E-05 |
| L-Tyrosine | 12.83656546 | 13.11856315 | 0.28199769 | 2.79369E-08 | 8.3252E-06 |
| Lysine | 13.52680435 | 13.94504356 | 0.418239204 | 2.46051E-05 | 0.006471134 |
| Malonic Acid | 19.14904933 | 20.67185962 | 1.522810289 | 2.04266E-14 | 6.51609E-12 |
| Maltose | 16.94730586 | 17.41940474 | 0.472098879 | 5.37272E-15 | 1.71927E-12 |
| Maltotriose | 16.22567427 | 17.49033613 | 1.264661867 | 1.30779E-35 | 4.45957E-33 |
| Mannitol | 19.74973922 | 21.62402957 | 1.874290353 | 9.14196E-46 | 3.16312E-43 |
| Meso-Erythritol | 18.57414641 | 19.86776769 | 1.293621287 | 1.0294E-53 | 3.59259E-51 |
| Methyldopa | 12.83238192 | 13.49934486 | 0.666962943 | 1.78352E-10 | 5.51109E-08 |
| N-Acetyl-D-Hexosamine | 17.38907379 | 17.8960245 | 0.50695071 | 1.36893E-19 | 4.53115E-17 |
| N-Propylamine | 17.66025419 | 17.12787266 | -0.532381529 | 5.58116E-07 | 0.000156272 |
| Ornithine | 16.59116371 | 17.23195676 | 0.640793042 | 0.000193989 | 0.048303338 |
| Palladium Diacetonate | 15.36692983 | 14.96965959 | -0.397270238 | 6.83832E-09 | 2.07201E-06 |
| Pelargonic Acid | 16.46545355 | 16.17271051 | -0.292743033 | 6.22521E-05 | 0.015998799 |
| Phosphoric Acid | 20.15238331 | 20.89515437 | 0.742771061 | 9.19099E-10 | 2.80325E-07 |
| Pipecolic Acid | 11.98049325 | 13.09685035 | 1.116357094 | 1.33297E-31 | 4.51878E-29 |
| Propyleneglycol | 16.55301572 | 15.74852988 | -0.804485841 | 2.03781E-07 | 5.84851E-05 |
| Pseudo Uridine | 15.82480091 | 16.44608365 | 0.621282742 | 4.82996E-14 | 1.53593E-11 |
| Pyrogallol | 11.51730934 | 13.96889719 | 2.451587844 | 1.0469E-07 | 3.0674E-05 |
| Quinic Acid | 13.40726078 | 14.51159984 | 1.104339066 | 6.67635E-30 | 2.25661E-27 |
| Ribitol | 19.5390554 | 20.03735797 | 0.498302575 | 3.36055E-13 | 1.06193E-10 |
| Ribonic Acid | 17.07450982 | 18.60141092 | 1.526901092 | 4.03095E-07 | 0.000113673 |
| Shikimic Acid | 14.49286862 | 15.22682741 | 0.733958783 | 9.2418E-09 | 2.79102E-06 |
| Sucrose | 17.23924642 | 18.7816232 | 1.542376783 | 1.10651E-16 | 3.60724E-14 |
| Tris(Trimethylsilyl)Methane | 14.46394861 | 15.21665976 | 0.752711148 | 2.33187E-08 | 6.9723E-06 |
| Urea | 23.05369933 | 23.59253366 | 0.538834337 | 9.48379E-08 | 2.78823E-05 |
| Uric Acid | 22.220133 | 23.32312536 | 1.102992354 | 7.60431E-44 | 2.62349E-41 |
| Xanthine | 12.76637973 | 13.8366192 | 1.070239463 | 3.27625E-07 | 9.30455E-05 |
| Xylitol | 16.42593994 | 17.85433329 | 1.428393352 | 4.18312E-39 | 1.43481E-36 |
| Z Hexose Pertms | 23.29508983 | 24.12857542 | 0.833485593 | 0.000104098 | 0.026336672 |
